# Protistan plankton communities in the Galápagos Archipelago respond to changes in deep water masses resulting from the 2015/16 El Niño

**DOI:** 10.1101/2021.05.06.441682

**Authors:** Erika F. Neave, Harvey Seim, Scott Gifford, Olivia Torano, Zackary I. Johnson, Diego Páez-Rosas, Adrian Marchetti

## Abstract

The Galápagos Archipelago lies within the eastern equatorial Pacific Ocean at the convergence of major ocean currents that are subject to changes in circulation. The nutrient-rich Equatorial Undercurrent upwells from the west onto the Galápagos platform, stimulating primary production, but this source of deep water weakens during El Niño events. From measurements collected on repeat cruises, the 2015/16 El Niño was associated with declines in phytoplankton biomass at most sites throughout the archipelago and reduced utilization of nitrate, particularly in large-sized phytoplankton in the western region. Protistan assemblages were identified by sequencing the V4 region of the 18S rRNA gene. Dinoflagellates, chlorophytes, and diatoms dominated most sites. Shifts in dinoflagellate communities were most apparent between the years; parasitic dinoflagellates, *Syndiniales*, were highly detected during the El Niño (2015) while the dinoflagellate genus, *Gyrodinium* dominated many sites during the neutral period (2016). Variations in protistan communities were most strongly correlated with changes in subthermocline water density. These findings indicate that marine protistan communities in this region are regimented by deep water mass sources and thus could be profoundly affected by altered ocean circulation.

## Introduction

The Galápagos Archipelago and surrounding waters (1-2 °S, 90-92 °W) are renowned for having diverse, highly productive ecosystems. The need to protect their marine ecosystems led to the establishment of the Galápagos Marine Reserve (GMR) in 1998 (Bensted-Smith, 1998). Despite greater efforts to conserve and study the GMR, one of the main sources of primary production, the marine protists, remain largely understudied. Protistan plankton exhibit diverse trophic modes, ranging from autotrophic phytoplankton to heterotrophic flagellates. A shift in protist community composition can therefore drastically alter the quantity of food available for higher trophic levels, and thus influence the overall productivity of the Galápagos marine ecosystem (De Vargas *et al*., 2015).

The convergence of ocean currents allows waters within the archipelago to have high primary production relative to the surrounding Eastern Equatorial Pacific (EEP) Ocean, a region considered high-nutrient, low-chlorophyll. The EEP is a high-nutrient, low chlorophyll region due to iron limitation (Behrenfeld *et al*., 1996), which is largely relieved in the GMR when currents upwell lithogenic nutrients from the Galápagos platform (Barber and Chavez, 1991; Rafter, Sigman and Mackey, 2017). Three major ocean currents provide the bulk of nutrient sources to the region (Lindley and Barber, 1998; Fiedler and Talley, 2006). The South Equatorial Current flows westward, enveloping the equator, and is fed by the Peruvian coastal upwelling and equatorial upwelling (Pennington *et al*., 2006) (Figure 1a). The North Equatorial Countercurrent flows eastward, transporting western Pacific warm pool water to the north of the South Equatorial Current. The most critical supply of nutrients to the region comes from the eastward-flowing Equatorial Undercurrent (EUC) which collides subsurface with the western side of the Galápagos platform. It flows around and through the archipelago below the surface layer (Kessler, 2006) carrying nutrient rich subtropical underwater, and can outcrop west of the archipelago, making this region a primary productivity hotspot (Chavez and Brusca, 1991; Sakamoto *et al*., 1998).

**Figure 1.**
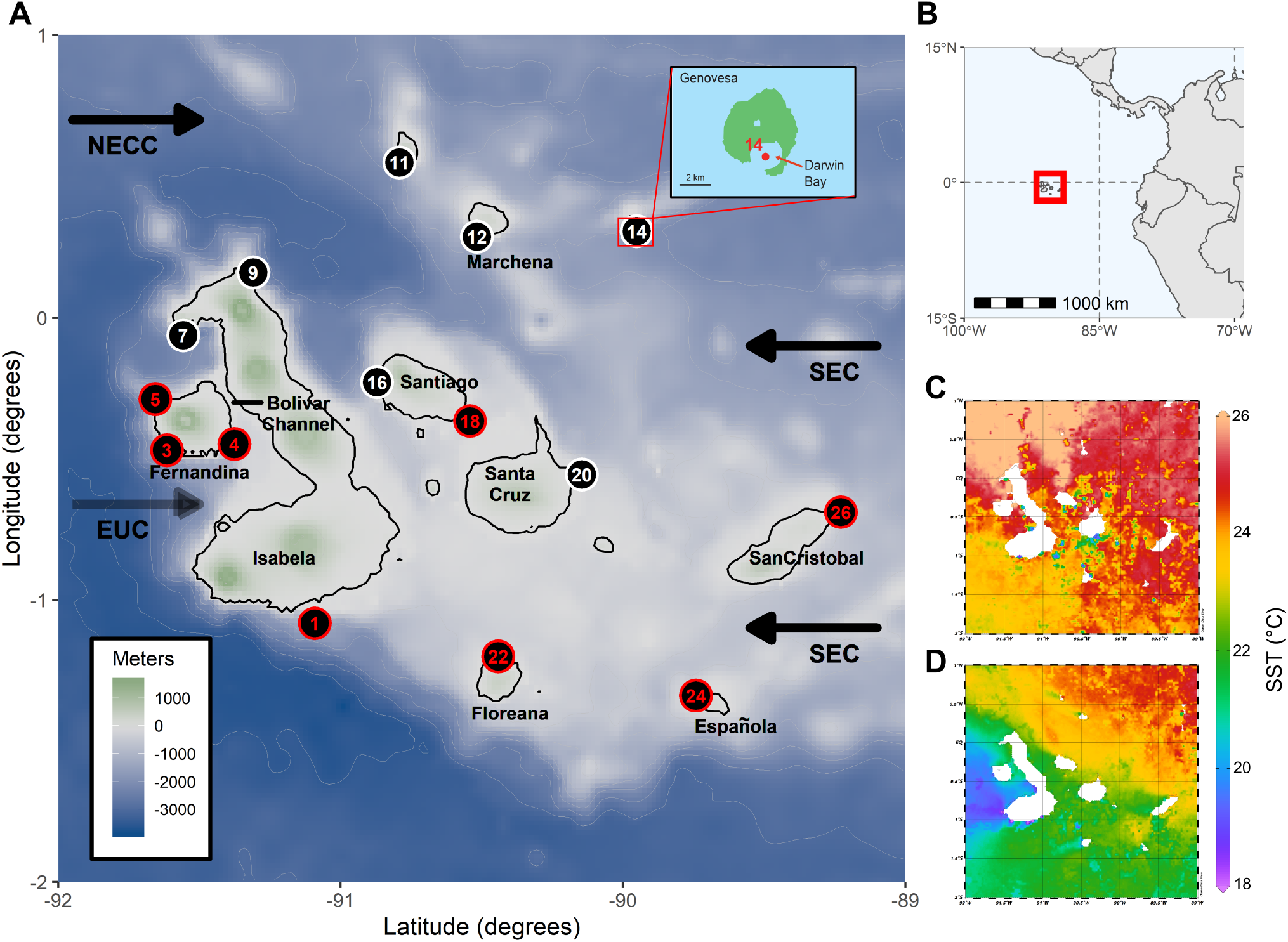
The Galápagos Archipelago study region. A) Topographic map of the Galápagos Archipelago and sampling locations. White numbers indicate sites which were sampled for oceanographic measurements while red numbers indicate sites that were sampled for oceanographic measurements and protistan community DNA. Currents are abbreviated such that: SEC = South Equatorial Current, NECC = North Equatorial Countercurrent, and EUC = Equatorial Undercurrent. Upper right inset shows the location of site 14 near Genovesa Island, a partially collapsed caldera. B) Map showing the location of the Galápagos Archipelago within the Eastern Equatorial Pacific. Monthly averaged sea surface temperatures (NOAA AVHRR) throughout the Galápagos Archipelago for the sampling period in C) October of 2015 and D) October of 2016.

Marine protistan communities in the EEP are influenced by El Niño Southern Oscillation (ENSO), a spatio-temporally complex interannual shift in equatorial Pacific Ocean circulation. The EEP harbors mostly small-sized phytoplankton communities consisting of *Prochlorococcus*, *Synechococcus*, and picoeukaryotes (Chavez *et al*., 1996). However, when equatorial upwelling is strong it can provide sufficient iron and silica for large-sized phytoplankton such as diatoms to bloom (Pennington *et al*., 2006; Masotti *et al*., 2010; Marchetti *et al*., 2010). During an El Niño event, EEP sea surface temperatures (SST) rise above the climatological average causing stratification. This results in weaker equatorial upwelling, a deeper thermocline, and a slower EUC. El Niño conditions lead to reduced nitrate availability and decreases in *Synechococcus*, likely because of their high cellular nitrogen requirement (Moore *et al*., 2002), which allow small heterotrophic protists to dominate, altering marine food web dynamics in the EEP (Masotti *et al*., 2010). Contrary to community dynamics in the EEP, on the west side of the Galápagos Archipelago, *Synechococcus* and *Prochlorococcus* concentrations decreased during a neutral period of stronger upwelling following the 2015/16 El Niño (Gifford *et al*., 2020). Given that *Prochlorococcus* is speculated to prefer nitrogen sources other than nitrate, it may be more competitive under stratified El Niño conditions (Moore *et al*., 2002). There is less understanding of how shifts in photosynthetic eukaryotes and other protists are influenced by ENSO in the Galápagos Archipelago specifically, where increased vertical mixing allows for a relatively higher baseline phytoplankton biomass than the surrounding EEP (Barber *et al*., 1996).

Despite limited knowledge on protistan communities, establishing causes for fluctuations in phytoplankton biomass in the GMR has been investigated previously (Maxwell, 1975; Feldman, 1986). Some studies have used satellite chlorophyll *a* (Chl *a*) proxies to understand phytoplankton biomass variability (Liu *et al*., 2014; Kislik *et al*., 2017). Despite seasonal Chl *a* variability in the GMR, its amplitude varies with basin-scale SST trends, in that Chl *a* peaks in the Boreal spring, synchronous with the strengthening of the EUC (Palacios, 2004; Sweet *et al*., 2007). However, this temporal pattern in Chl *a* amplitude does not hold true spatially, as individual bioregions of the GMR differ (Edgar *et al*., 2004). The South Equatorial Current and the North Equatorial Countercurrent meet forming the equatorial front. The seasonal oscillation of this has been used to predict patterns of Chl *a* concentration (Schaeffer *et al*., 2008). In the latter part of the year, the strengthening and tilt of the equatorial front can largely explain Chl *a* variability (Palacios, 2004).

Fewer studies have focused on observing in situ environmental conditions and their influence on protistan community compositional changes (McCulloch, 2011; Carnicer *et al*., 2019). These observations are necessary for understanding the ecological implications of El Niño events in this region. Sporadic surveys of phytoplankton communities provide a historical record of common diatoms and dinoflagellate species, however they are limited to observations of large, more “charismatic” species easily identified using light microscopy (Torres, 1998; Torres and Tapia, 2000, 2002; Tapia and Naranjo, 2012). Some harmful algal species were identified along with spatial variability in dinoflagellate diversity, which was attributed to changes in deep water masses to the west of the Galápagos platform (Carnicer *et al*., 2019). Accessory phytoplankton pigments have also been used to assess phytoplankton composition in the GMR, such that relative abundances of diatoms and chlorophytes were found to decrease during the 2004/05 El Niño event, while cyanobacteria and haptophyte proportions increased (McCulloch, 2011).

In this study, DNA metabarcoding (i.e., sequencing the 18S rRNA [18S] gene) was used to examine protistan community composition. Here we show distinct shifts in protist plankton genera in waters of the Galápagos Archipelago and how they correlate with primary productivity and oceanographic variables during the 2015/16 ENSO cycle. Because protistan plankton are important links between oceanographic processes and marine food webs, it is imperative to understand factors influencing their composition and production with shifts in environmental conditions attributed to climatic events.

## Methods and materials

### Sample collection

Fifteen sites within the GMR (89 – 92 °W, 1.5 °S – 2 °N) were sampled over October 10^th^ to 24^th^ of 2015 and October 19^th^ to November 11^th^ of 2016 (Figure 1a). Based on the Oceanic Niño Index and the location of the GMR, situated within both the Niño 1.2 region (80 – 90 °W, 0 – 10 °S) and the Niño 3 region (150 – 90 °W, 5 °N – 5 °S), our sampling periods occurred during El Niño (2015) and neutral conditions (2016) (Santoso, Mcphaden and Cai, 2017). Remote sensing of sea surface temperatures in the region indicate significantly warmer surface waters during the sampling period in 2015 relative to 2016 (Figure 1c-d). Using a Seabird SBE 19plus V2 SeaCAT Profiler, CTD profiles of temperature, salinity and photosynthetically active radiation (PAR) were collected to ∼100 m depth. Ten liter Niskin bottles were used to collect seawater from the euphotic zone at 50%, 30%, 10%, and 1% incident irradiance (I_o_) depths to measure dissolved inorganic nutrients, Chl *a*, dissolved inorganic carbon (DIC) and nitrate (NO_3_^−^) uptake rates, and picoplankton cell counts. Seawater was dispensed into acid-cleaned, seawater rinsed 10 L Cubitainers (Hedwin Corporation, Newark, DE, USA) and subsampled for measurements. Additional seawater was collected from 50% I_o_ to obtain DNA (Figure 1a). Not all sites yielded DNA concentrations sufficient for sequencing. Processed filters and samples were frozen at −20 °C until onshore analysis.

### Seawater Properties

Temperature and salinity profiles were used to determine the depths of the mixed and subthermocline layers. All CTD casts were corrected using SeaBird’s SeaSoft software. Potential density (hereafter referred to as ‘density’) was calculated using the sw_pden() function from the Mixing Oceanographic toolbox v 1.8.0.0 in MATLAB (R2017b). A consistent density structure was observed, of a surface and deep layer of almost uniform density, separated by a density gradient (interfacial layer) that was typically 10s of meters thick. The surface mixed layer depth was defined as the depth at which the change in density from the surface was > 0.35 kg/m^3^. The subthermocline layer depth (the top of the deep layer of uniform density) was determined by calculating the depth at which change in density from the bottom of the cast was > 0.2 kg/m^3^. The casts were visually inspected to ensure that the density cut-off values defined the layers appropriately. Temperature, salinity, and density of the mixed and subthermocline layers were averaged from CTD cast measurements. Methods for measuring dissolved inorganic nutrients are described in the supplemental material.

### Phytoplankton Biomass and Productivity

Phytoplankton biomass was approximated by measuring size-fractioned (<5 μm and >5 μm) Chl *a* concentrations using the acetone extraction method (Sartory and Grobbelaar, 1984). Picophytoplankton cells, specifically *Prochlorococcus*, *Synechococcus*, and picoeukaryote populations were enumerated using flow cytometry (Johnson *et al*., 2010). Size-fractionated DIC (i.e. primary productivity) and NO_3_^−^ uptake rates were measured from 24 hr incubations using stable isotope tracer techniques (Dugdale and Goering, 1967; Hama *et al*., 1983). Additional methods describing Chl *a*, flow cytometry, primary productivity and NO_3_^−^ uptake measurements are provided in the supplemental material.

### DNA sequencing and Bioinformatics

Protistan taxonomic identification and proportions were obtained through sequencing the V4 region of the 18S rRNA gene. DNA collection and amplicon library preparation are described in the supplemental material. Approximately four million paired-end reads were obtained using the Illumina MiSeq sequencing platform across two lanes. Mean amplicon length for sequencing lane one was 561 bp, while mean amplicon length for sequencing lane two was 599 bp. Sequence files were demultiplexed. QIIME 2 v.2018.6 software was used for processing the raw reads into assembled amplicons. The QIIME 2 plug-in, DADA2 was used for denoising such that reverse and forward reads were truncated to 250 base pairs (bp) and 210 bp, and 260 bp and 280 bp, for sequencing lanes one and two respectively. Chimeras were removed by the consensus method and reads were merged (Supplementary Table 1) using default scripts provided in the QIIME 2 documentation (docs.qiime2.org). Assembled amplicons were annotated by a BLAST search against the SILVA v. 123 reference database using a 90% pairwise identity cutoff. Metazoans were removed. Using the R package phyloseq v. 1.24.2, the samples were rarefied to 2066 reads per site (Supplementary Figure 1).

Annotations which were unknown in the highest taxonomic ranks (Kingdom, Phylum, Class, Order) were removed, under the assumption that it was unlikely to sample a novel high taxonomic rank of plankton. Custom taxonomy was assigned to the Class taxonomic rank based on the top seven relatively abundant groupings (Supplementary Table 2). Any annotation that did not belong to the top seven groups was annotated as “Other”. Unknown or uncultured annotations in the lowest three taxonomic ranks (Family, Genus, Species) were grouped into the “Other” category within their respective Class ranks. Additionally, where possible, the top seven genera were maintained while the rest were also annotated as “Other” (there were only six known genera in the Chlorophyta). This resulted in 120 genera, prior to grouping unknown and uncultured OTUs (Supplementary table 2). All raw sequences have been deposited in the NCBI SRA database (Accession No. PRJNA689599).

### Statistical Analyses

Analysis of water properties, phytoplankton biomass, productivity, and 18S data was performed using the vegan package (v. 2.5-4) in R version 3.6.2. Summary statistics and Mann-Whitney-Wilcoxon tests were used to test for differences in dissolved nutrients, Chl *a*, primary productivity and NO_3_^−^ uptake rates between the two sampling years (Supplementary Table 3). The 18S data was arranged in an OTU table and transformed to relative proportions at the genus level, from which Bray-Curtis dissimilarity distances were calculated using the vegdist() function. All other variables (i.e., physical water properties, phytoplankton biomass, primary productivity and NO_3_^−^ uptake rates) were similarly transformed to Euclidean distances. A correlation matrix was used to assess which variables were relatively redundant (Supplementary Figure 2). These variables were removed for the BIO-ENV (‘biota-environment’) analysis. The bioenv() function was used to find the best subset of variables which had the maximum rank correlation with the community Bray-Curtis dissimilarities (Clarke and Ainsworth, 1993). These variable subsets were identified for the entire protistan community and for the following subgroups of the community: dinoflagellates, chlorophytes, and diatoms. A Non-metric Multi-Dimensional Scaling (NMDS) plot was made from the Bray Curtis community dissimilarity distances. The envfit() function was then used to fit the best subsets of variables onto the community dissimilarity ordinations (Supplementary Table 4).

## Results and Discussion

### Physical seawater properties

Differences in upper ocean physical seawater properties between the El Niño (2015) and neutral (2016) years were apparent (Figure 2, Supplementary Table 5). At most sites, mixed layer depths were deeper in 2015 then 2016 (Fig. 2a). The mixed layer temperature range was warmer in 2015 (22.9-26.4 °C) relative to 2016 (17.1-23.7 °C), with a mean difference of 3.8°C (Figure 2b). Mixed layer salinities varied by an average of 0.23 PSU, with most being higher in 2015 (Figure 2c). As a result, the mean density in 2016 was higher by 0.99 kg/m^3^ during the neutral period (Figure 2d). These differences in densities also existed spatially, such that the sites located west of Isabela Island (Sites 3-7) had the highest mixed layer density values in 2016 (Figure 2d). Similar interannual density trends were observed in the subthermocline layer, where subthermocline depths at most sites were deeper in 2015 than 2016 (Fig. 2e). The subthermocline layer was on average 3.2 °C cooler and 0.07 PSU fresher in 2016 resulting in seawater which was on average 0.70 kg/m^3^ denser relative to the El Niño (Fig. 2f-h). One notable exception was site 14 located near Genovesa Island, inside a partially submerged caldera (see inset map in Fig. 1a). The caldera is isolated from surrounding ocean by an approximately 10 m deep sill, restricting exchange with waters outside the caldera and allowing seawater properties within the caldera to remain more constant between years. Aside from this site outlier, subthermocline layer densities showed consistent temporal change relative to the mixed layer, which was more spatially sensitive.

**Figure 2.**
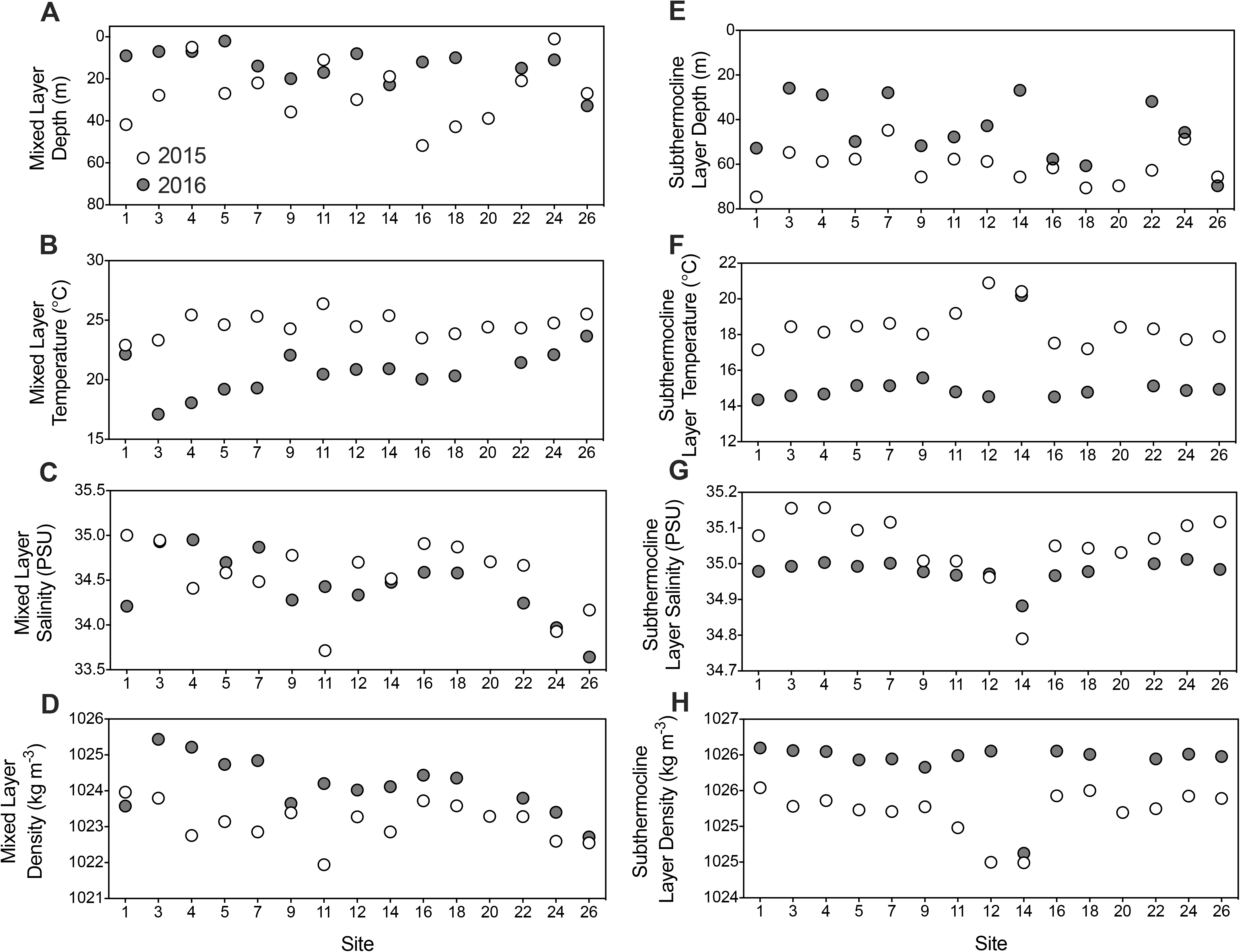
Physical seawater properties during the 2015 and 2016 sampling period. A) Mixed layer depths, B) mixed layer temperatures, C) mixed layer salinities, D) mixed layer densities, E) subthermocline layer depths, F) subthermocline layer temperatures, G) subthermocline layer salinities and H) subthermocline layer densities.

The differences in water mass densities between the years were reflected in dissolved nutrient concentrations. Nitrate and silicic acid (Si[OH]_4_) concentrations within the mixed layer were lower at nearly all sites in 2015 relative to 2016 (Fig. 3a and b). Nitrate concentration in the euphotic zone was significantly lower during the El Niño relative to 2016, with median values of 1.70 µmol L^−1^ and 6.20 µmol L^−1^ respectively (Fig. 3c). Similarly, Si(OH)_4_ had lower median values in 2015 (1.65 µmol L^−1^) compared to 2016 (4.99 µmol L^−1^) (Fig. 3d). Both years had higher NO_3_^−^ concentrations relative to Si(OH)_4_ concentrations, however the Si(OH)_4_: NO_3_^−^ slope of the distribution in 2015 was lower (m = 0.40) than in 2016 (m = 0.69), indicating that the concentration of Si(OH)_4_ relative to NO_3_^−^ was lower during the El Niño (Figure 3e). Overall, physical seawater properties and nutrient concentrations in the GMR show strong temporal trends indicative of oceanographic variation during the 2015/16 El Niño event.

**Figure 3.**
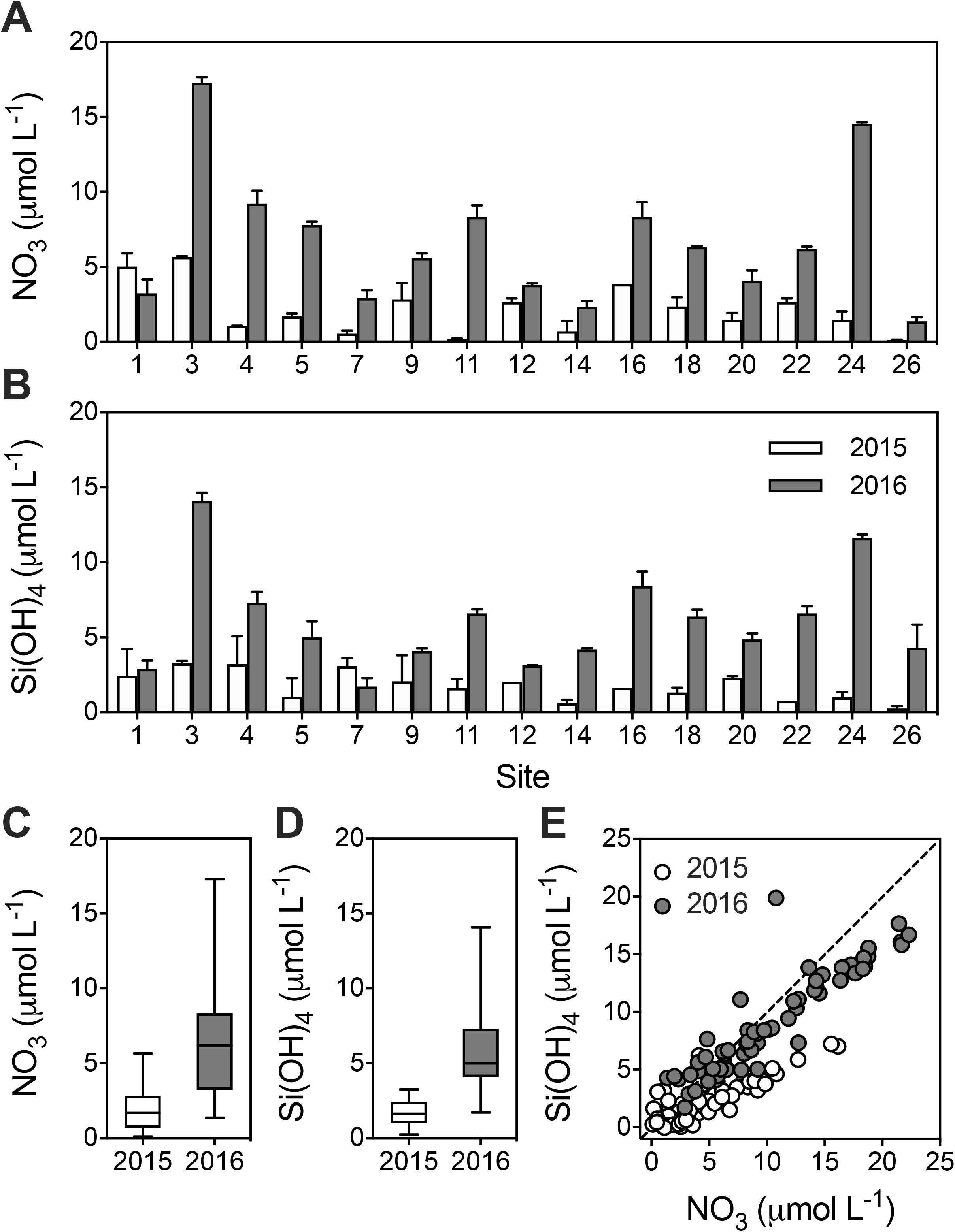
Dissolved inorganic nutrients during the 2015 and 2016 sampling period. A) Nitrate (NO_3_^−^) concentrations. B) silicic acid (Si[OH]_4_) concentrations. C) interquartile range of NO_3_ concentrations at 50% incident irradiance. D) interquartile range of Si(OH)_4_ concentrations at 50% incident irradiance. E) Scatter plot of NO_3_^−^ verses Si(OH)_4_ concentrations collected at depths (50%, 30%, 10%, 1% incident irradiance) throughout the euphotic zone. The dashed line represents the 1:1 line. Error bars indicate the standard deviation of the mean (n=2).

### Phytoplankton biomass and primary productivity

Chl *a* concentrations were higher in 2016 at most sites with the primary exceptions of sites 14, and 16 (Fig. 4a). The median concentration of the small-size fraction (< 5 μm) was similar between 2015 (0.22 µg L^−1^) and 2016 (0.23 µg L^−1^) (Fig. 4b). The large-size fraction (> 5 μm) had a higher median in 2016 (0.2 µg L^−1^) relative to 2015 (0.13 µg L^−1^) with a maximum concentration of 1.43 µg L^−1^ in 2016 (Fig. 4c). These are comparable to measurements (89 – 92°W, 2 °S – 1°N) observed previously during a neutral period, where the average surface Chl *a* was 0.25 µg L^−1^ and the highest concentration (0.53 µg L^−1^) was recorded west of Isabela island (Torres and Tapia, 2000). In our study, the average small and large size-fractions of Chl *a* did not significantly differ between years, however the large size-fraction showed stronger temporal (*p*=0.06) and spatial trends, having higher concentrations in 2016 at sites located west of Isabela Island (Fig. 4a and 4d).

**Figure 4.**
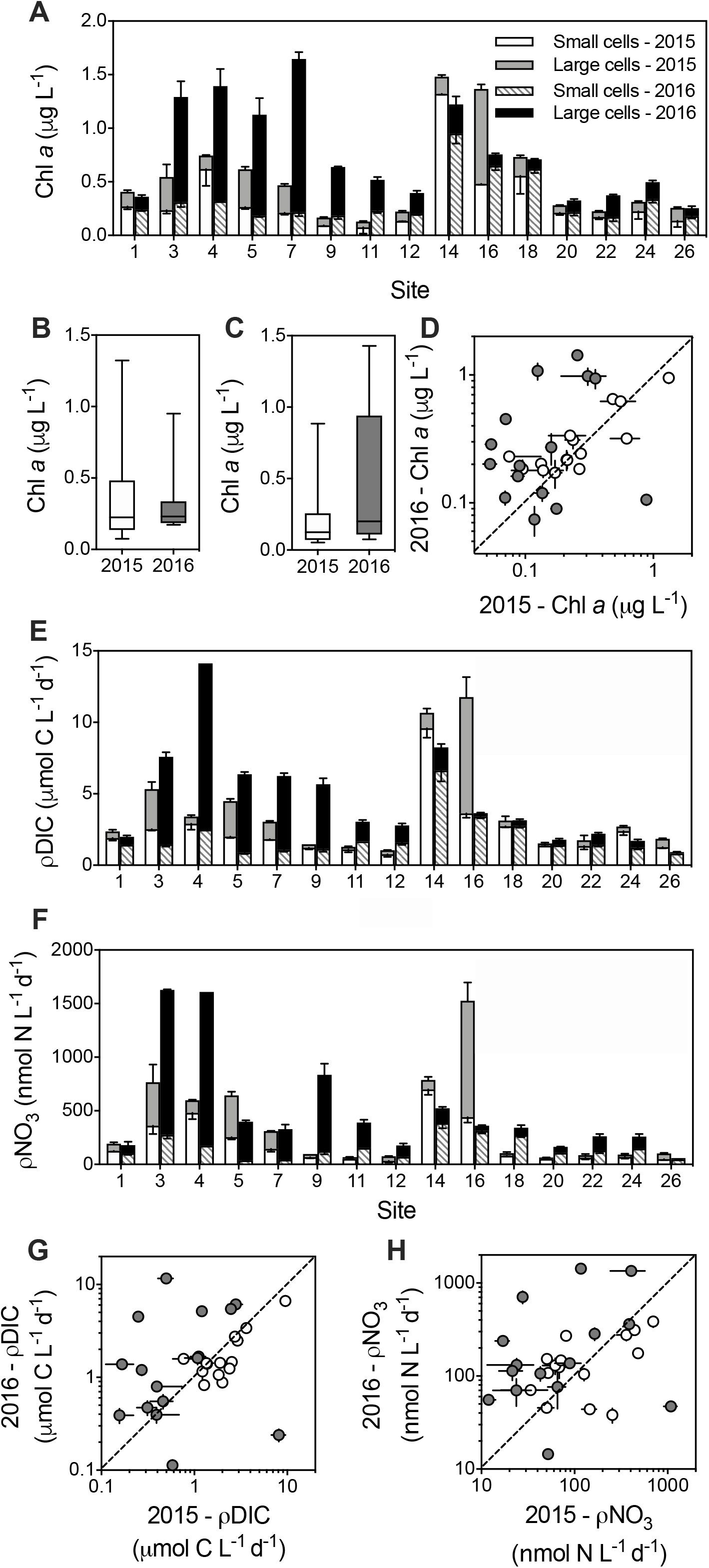
Phytoplankton biomass and primary productivity at 50% incident irradiance during the 2015 and 2016 sampling period. A) Size-fractionated chlorophyll *a* (Chl *a*) concentrations. Chl *a* is distinguished between large (> 5 μm) and small (<5 μm) size-fractions. B) interquartile range of Chl *a* concentrations in small cells across all sites and C) interquartile range of Chl *a* concentrations in large cells across all sites. D) Scatter plot of large (grey) and small (white) cell Chl *a* concentrations in 2016 versus 2015. Size-fractionated E) dissolved inorganic carbon uptake rates (i.e. primary productivity; ρDIC) and F) NO_3_^−^ uptake rates (ρNO_3_^−^). Bar colors are the same as in A. Scatter plot of size-fractionated G) ρDIC and H) ρNO_3_^−^ in 2016 versus 2015. The dashed lines represent the 1:1 line. Symbol colors are the same as in D. Error bars indicate the standard deviation of the mean (n=3). In A, E and F, error bars for the small size-fraction are in the downward direction whereas error bars for the large size-fraction are in the upward direction.

*Synechococcus* flow cytometry cell counts were higher at western sites 3-7 in 2015 than 2016 (Supplementary Figure 3), similar to observations made at the same sites quantified using metagenomics (Gifford *et al*., 2020). Site four had the greatest difference between the years, having a concentration of 2.4×10^5^ cells mL^−1^ in 2015 and 5.9 ×10^4^ cells mL^−1^ in 2016. The highest *Synechococcus* concentration (2.6×10^5^ cells mL^−1^) was recorded in 2015 at site 14, within the isolated caldera. *Prochlorococcus* was generally more abundant in 2015, particularly in the central and eastern sites, and ranged in concentration from 6.2 ×10^3^ cells mL^−1^ to 1.9×10^5^ cells mL^−1^ (Supplementary Figure 3). In 2016 *Prochlorococcus* concentrations were generally lower, ranging from 3.8×10^3^ cells mL^−1^ to 1.6×10^4^ cells mL^−1^ except at sites five and seven where they were an order of magnitude lower. Picoeukaryotes typically ranged from 10^3^-10^4^ cells mL^−1^ and had more complex abundance patterns between years than *Synechococcus* and *Prochlorococcus* (Supplementary Figure 3). For example, there were higher concentrations of picoeukaryotes at western sites 4-7 in 2015, while lower concentrations relative to 2016 were recorded at sites 20-26.

Total DIC uptake rates (primary productivity) remained commensurate between years but showed spatial variability, such that sites 3-12 and 18-22 increased during 2016 while other sites decreased (Figure 4e). Insignificant temporal trends in total primary productivity could be attributed to opposite trends in the phytoplankton size-fractions. For instance, median DIC uptake rates in the large size-fraction were 0.46 µmol C L d^−1^ in 2015 to 1.2 µmol C L d^−1^ in 2016, while the small cells decreased from having a median of 1.9 µmol C L d^−1^ in 2015 to 1.44 µmol C L d^−1^ in 2016 (Figure 4g). NO_3_^−^ uptake rates displayed trends similar to Chl *a* and primary productivity such that the large size-fraction differed more between years than the small size-fraction (*p*=0.07; *p*=0.87) (Figure 4f and h). The decrease in biomass of the large size-fraction in 2015 coincided with a lower median NO_3_^−^ uptake rate (51.9 nmol N L d^−1^) in the large size-fraction compared to the small size-fraction (81.4 nmol N L d^−1^) (Figure 4h). Overall median rates of NO_3_^−^ uptake were greater in both size-fractions during 2016 (129.8, 131.6 nmol N L d^−1^; small and large size-fractions, respectively) (Figure 4h).

### Shifts in protistan community composition

The most proportionally dominant protist groups in the GMR included the dinoflagellates (part of the Alveolata group), chlorophytes, diatoms (part of the Stramenopiles group), Hacrobia, Opisthokont fungi, and Rhizaria (Figure 5a). Collectively, dinoflagellates, chlorophytes, and diatoms dominated the protistan communities and exhibited the most variability between years (Figure 5b-d). Other groups had high spatial variability across sites. For instance, the ciliates and rhizarians were detected at every site in 2015 but lacked detection or genus level identification at over half of the sites in 2016 (Supplemental Fig. 3). Thus, analyses were focused on changes in the three most dominant groups.

**Figure 5.**
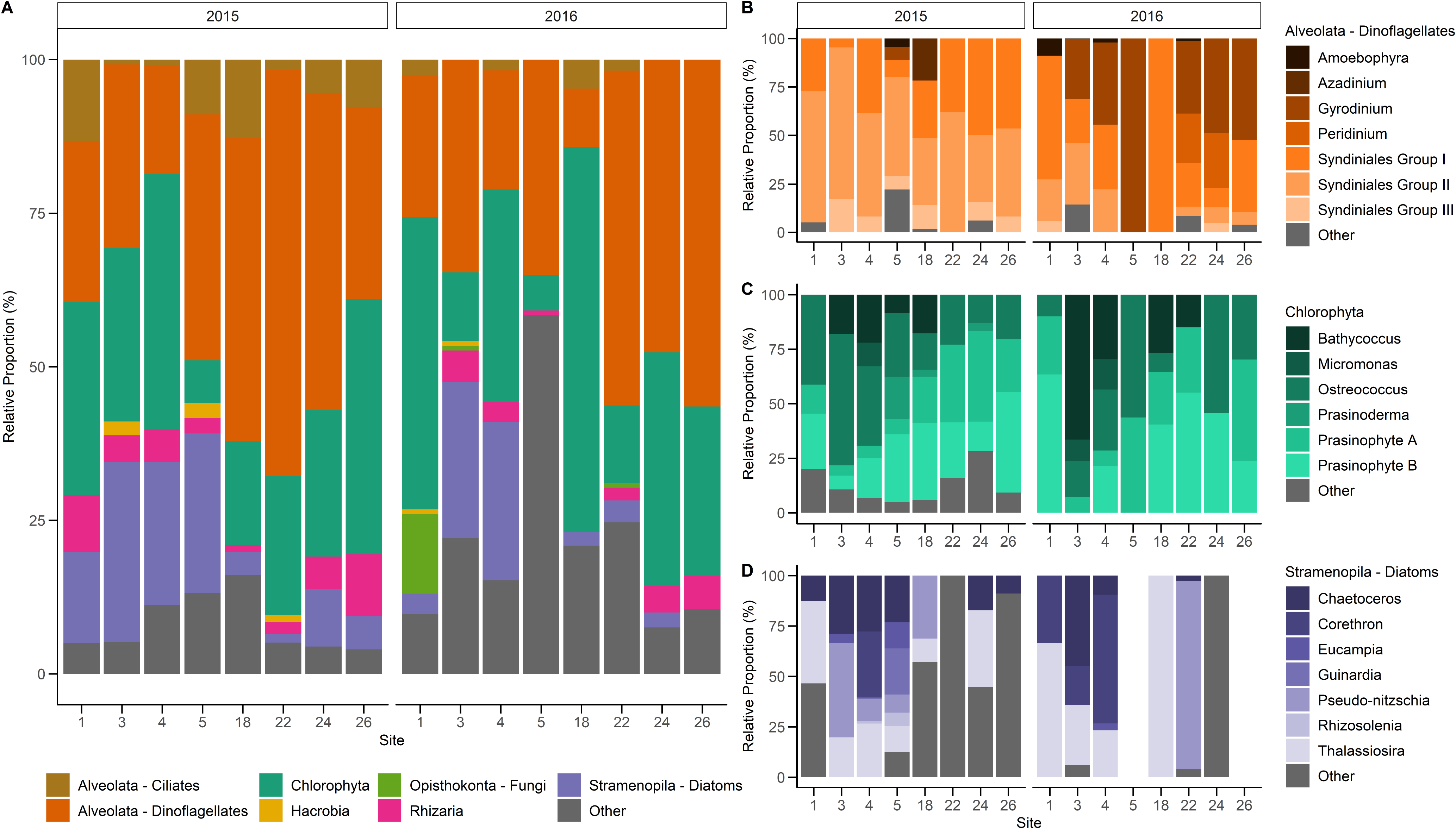
Protistan community composition based on 18S rRNA gene amplicons. A) Relative proportions of protists at class level groupings in 2015 and 2016. B) Relative proportions of the Dinoflagellate group, highlighting the top seven most abundant genera. C) Relative proportions of the Chlorophyta group, highlighting the top six most abundant genera. D) Relative proportions of the Diatom group, highlighting the top seven most abundant genera.

The dinoflagellates had a total of 186 OTUs within 21 genera. Dinoflagellate communities were proportionally well represented by seven primary order and genus level groups (Figure 5b). Temporal changes in dinoflagellate proportions between years were more prominent than spatial changes among sites, as indicated by a larger variation along the horizontal axis of the NMDS plot (Figure 6). Changes in dinoflagellate community structure most resembled the collective changes in the density of the subthermocline layer, primary productivity by small cells, and picoeukaryote cell abundance (Spearman’s rho = 0.61) (Table 1). Dinoflagellate composition changed the most between years relative to the chlorophytes and diatoms, yet as a broader group they maintained relatively high proportions over the 2015/16 ENSO. This could be due in part to the diverse life strategies that enable dinoflagellates to bloom at various phases of the upwelling cycle (Smayda and Trainer, 2010).

**Figure 6.**
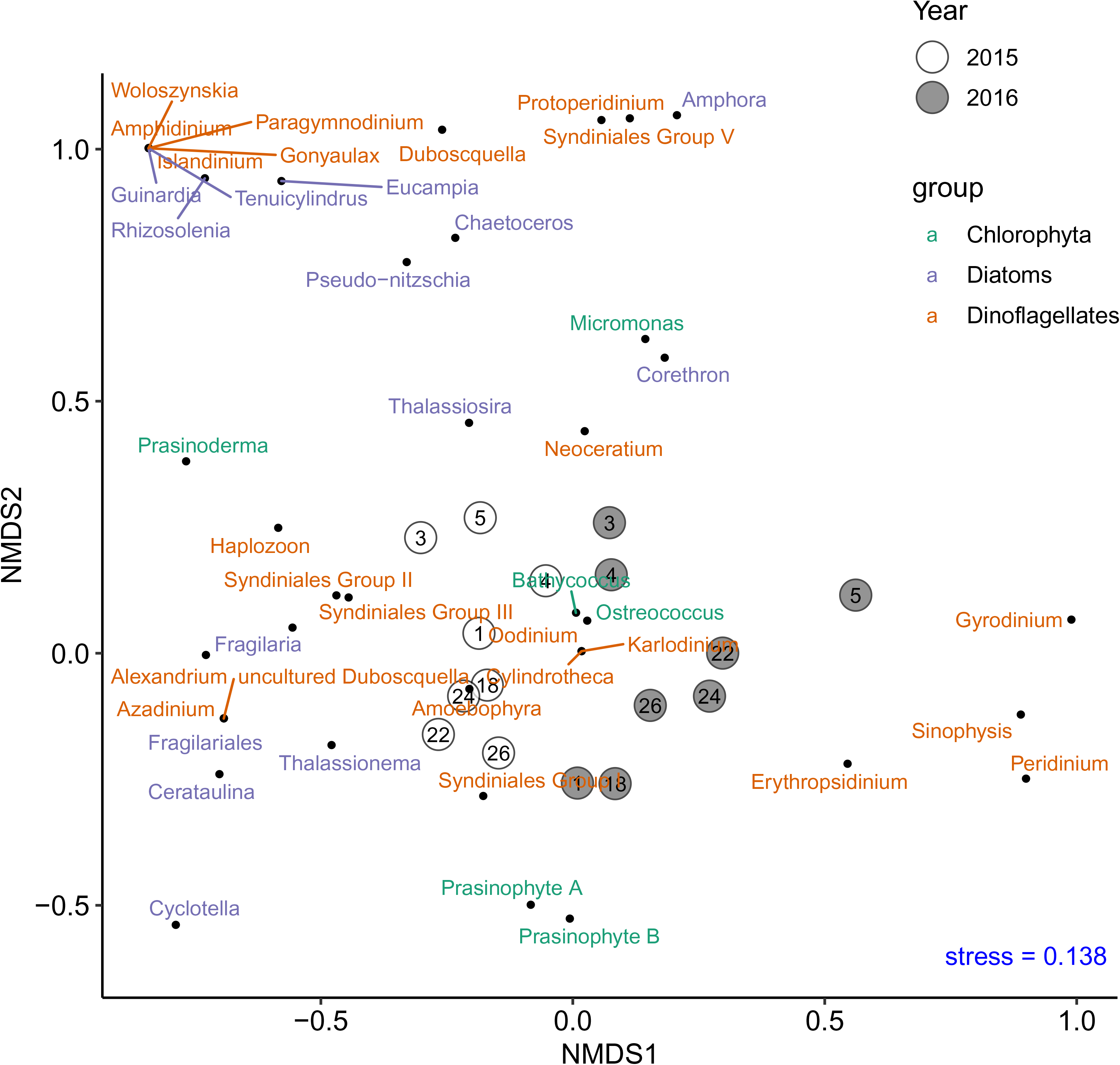
Non-metric Multi-Dimensional Scaling (NMDS) plot showing differences in the whole protistan community composition based on the Bray-Curtis dissimilarity matrix. 2015 sites (white) and 2016 sites (grey) are labelled with site numbers. The loadings for each genera group are plotted as points (black).

**Table 1.** Combinations of variables yielding the ‘best matches’ of variable (Euclidian) and community (Bray Curtis dissimilarity) matrices using Spearman’s rank correlation (rho). The variables are abbreviated such that: lg_updic = dissolved inorganic carbon uptake by the large (>5 µm) phytoplankton size-fraction; pden_dl = potential density of the subthermocline layer; phosphate = phosphate concentration; pico = picoeukaryote flow-cytometry counts; pro = *Prochlorococcus* flow-cytometry counts; sil = silicic acid concentration; sm_updic = dissolved inorganic carbon uptake by the small (<5 µm) phytoplankton size-fraction; sm_upnit = nitrate uptake by the small (<5 µm) phytoplankton size-fraction; syn = *Synechococcus* flow-cytometry counts; temp_ml = temperature of the mixed layer; tot_upnit = total nitrate uptake of both large and small phytoplankton size-fractions. (a) Whole protist community (b) Dinoflagellates (c) Chlorophytes (d) Diatoms

|  | Environmental & Ecological Variables | Variables | Spearman's<br>Rho |
| --- | --- | --- | --- |
| **A. Whole Community** |  |  |  |
|  | tot_upnit, pico, pro, pden_dl | 4 | 0.6239 |
|  | tot_upnit, pico, pden_dl | 3 | 0.6216 |
|  | tot_upnit, syn, pico, pro, pden_dl | 5 | 0.6189 |
|  | tot_upnit, sm_updic, syn, pico, pro, pden_dl | 6 | 0.6179 |
|  | sil, tot_upnit, sm_updic, syn, pico, pro, pden_dl | 7 | 0.6099 |
|  | sil, sm_upnit, tot_upnit, sm_updic, syn, pico, pro, pden_dl | 8 | 0.6057 |
|  | sil, sm_upnit, tot_upnit, sm_updic, syn, pico, pro,<br>pden_dl, temp_ml | 9 | 0.5898 |
|  | sil, sm_upnit, tot_upnit, sm_updic, lg_updic, syn, pico,<br>pro, pden_dl, temp_ml | 10 | 0.5676 |
|  | syn, pden_dl | 2 | 0.5664 |
|  | phosphate, sil, sm_upnit, tot_upnit, sm_updic, lg_updic,<br>syn, pico, pro, pden_dl, temp_ml | 11 | 0.5498 |
|  | pden_dl | 1 | 0.3681 |
| **B. Dinoflagellates** |  |  |  |
|  | sm_updic, pico, pden_dl | 3 | 0.6063 |
|  | sm_updic, pico, pro, pden_dl | 4 | 0.5951 |
|  | pico, pden_dl | 2 | 0.566 |
|  | sil, sm_updic, syn, pico, pden_dl | 5 | 0.5649 |
|  | sil, sm_updic, syn, pico, pro, pden_dl | 6 | 0.5495 |
|  | sil, sm_updic, syn, pico, pro, pden_dl, temp_ml | 7 | 0.5183 |
|  | phosphate, sil, sm_updic, syn, pico, pro, pden_dl,<br>temp_ml | 8 | 0.4965 |
|  | phosphate, sil, sm_updic, lg_updic, syn, pico, pro,<br>pden_dl, temp_ml | 9 | 0.4748 |
|  | phosphate, sil, sm_upnit, sm_updic, lg_updic, syn, pico,<br>pro, pden_dl, temp_ml | 10 | 0.4493 |
|  | pden_dl | 1 | 0.4222 |
|  | phosphate, sil, sm_upnit, tot_upnit, sm_updic, lg_updic,<br>syn, pico, pro, pden_dl, temp_ml | 11 | 0.4143 |
| **C. Chlorophytes** |  |  |  |
|  | tot_upnit, pico | 2 | 0.6826 |
|  | sm_upnit, syn, pico | 3 | 0.6756 |
|  | sm_upnit, syn, pico, pden_dl | 4 | 0.6739 |
|  | sm_upnit, tot_upnit, syn, pico, pden_dl | 5 | 0.6733 |
|  | sm_upnit, tot_upnit, syn, pico, pro, pden_dl | 6 | 0.6621 |
|  | sm_upnit, tot_upnit, lg_updic, syn, pico, pro, pden_dl | 7 | 0.6315 |
|  | sm_upnit, tot_upnit, lg_updic, syn, pico, pro, pden_dl,<br>temp_ml | 8 | 0.5947 |
|  | pico | 1 | 0.5861 |
|  | phosphate, sm_upnit, tot_upnit, lg_updic, syn, pico, pro,<br>pden_dl, temp_ml | 9 | 0.5655 |
|  | phosphate, sm_upnit, tot_upnit, sm_updic, lg_updic, syn,<br>pico, pro, pden_dl, temp_ml | 10 | 0.5381 |
|  | phosphate, sil, sm_upnit, tot_upnit, sm_updic, lg_updic,<br>syn, pico, pro, pden_dl, temp_ml | 11 | 0.4993 |
| **D. Diatoms** |  |  |  |
|  | sm_updic, pro | 2 | 0.3671 |
|  | sil, sm_updic, pro | 3 | 0.3585 |
|  | sm_updic | 1 | 0.3353 |
|  | sil, tot_upnit, sm_updic, pro | 4 | 0.3312 |
|  | sil, sm_upnit, tot_upnit, sm_updic, pro | 5 | 0.2969 |
|  | sil, sm_upnit, tot_upnit, sm_updic, pro, pden_dl | 6 | 0.2788 |
|  | sil, sm_upnit, tot_upnit, sm_updic, pico, pro, pden_dl | 7 | 0.2554 |
|  | sil, sm_upnit, tot_upnit, sm_updic, lg_updic, pico, pro,<br>pden_dl | 8 | 0.2341 |
|  | sil, sm_upnit, tot_upnit, sm_updic, lg_updic, syn, pico,<br>pro, pden_dl | 9 | 0.2017 |
|  | phosphate, sil, sm_upnit, tot_upnit, sm_updic, lg_updic,<br>syn, pico, pro, pden_dl | 10 | 0.1726 |
|  | phosphate, sil, sm_upnit, tot_upnit, sm_updic, lg_updic,<br>syn, pico, pro, pden_dl, temp_ml | 11 | 0.1444 |

Members of the dinoflagellate genus *Gyrodinium* showed the strongest temporal shift, seeming to favor the relatively cooler, higher nutrient conditions present in 2016 such that they were notably represented in the west (sites 3-5) and east (sites 22-26) (Figure 4e). Some *Gyrodinium* species have been found in sediments, and are suspected to form benthic resting cysts which live on internal nutrient reserves for long periods of dormancy until conditions become more favorable for growth (Shang *et al*., 2019). Along the coasts of Santa Cruz and other small neighboring islands in the central region of the GMR, water column surveys of dinoflagellate communities found that 84% of samples contained benthic epiphytic dinoflagellates (Carnicer *et al*., 2019), a high percentage given that only ∼10% of dinoflagellates associate with a substrate (Hoppenrath *et al*., no date). The suspension of these epiphytic dinoflagellates in the GMR suggests potential physical mechanisms for benthic resting cyst resuspension. Other cyst forming genera which were detected in our samples include *Alexandrium*, *Gonyaulax*, *Neoceratium*, *Paragymnodinium*, *Peridinium*, *Protoperidinium*, and *Woloszynskia* (Pospelova and Head, 2002; Bravo and Figueroa, 2014; Yokouchi, Onuma and Horiguchi, 2018). *Peridinium* was also detected in 2016 at sites 22 and 24, where it made up a significant proportion of the dinoflagellate community (Figure 5b). Long-term nutrient and temperature stress are the most common causes of resting cyst formation in dinoflagellates (Bravo and Figueroa, 2014). For example, *Gyrodinium uncatenum,* now renamed *Levanderina fissa*, forms cysts to survive in a dormant resting stage which can last for a duration of months to decades (Anderson, Coats and Tyler, 1985). This and other direct observations of cyst formation in closely related species, suggest that the *Gyrodinium* genus have the ability to survive long-term environmental stress (Bravo and Figueroa, 2014; Shang *et al*., 2019). Therefore, resting cyst formation may be an important strategy for dinoflagellates to subsist over El Niño events.

Syndiniales are a group of parasitoid dinoflagellates that survive via dinoflagellate or metazoan hosts (Jephcott *et al*., 2016). Metabarcoding techniques have revealed that these parasites are more prevalent than previously recognized (Guillou *et al*., 2008), and this too is the case in the GMR. Syndiniales groups I and II were dominant in 2015; both groups were simultaneously detected in 2016 except at sites five and 18. In 2016, site 18 notably had a dinoflagellate community highly dominated by Syndiniales I. Syndiniales III covered a larger spatial extent in 2015 than 2016 but generally followed the same trend as Syndiniales I and II, such that relative proportions were typically less in 2016 (Figure 5b). One caveat of our metabarcoding approach is that it does not distinguish between free-living cells and host associated parasites, which makes the ecological role of Syndiniales difficult to assess. Notwithstanding, higher relative proportions of Syndiniales during the El Niño may be a result of increased infection rates since host cell death precedes the release of Syndiniales spores (Jephcott *et al*., 2016; Clarke *et al*., 2019). *Ameobophyra*, a specific Syndiniales genus identified in the GMR (Figure 5b), is estimated to use half of its host’s biomass for spores leaving the other half as particulate and dissolved organic matter (Salomon and Stolte, 2010; Jephcott *et al*., 2016). While Syndiniales can exploit photosynthetic hosts, Syndiniales I have been found to correlate positively with Chl *a*, perhaps implying its association with high host biomass or productive regions (Clarke *et al*., 2019). Despite the high detection of Syndiniales, particularly during the El Niño, their effect on primary productivity and food web dynamics remains unclear.

The chlorophyta or green algae had a total of 163 identified OTUs within six genera. The six genera are displayed with an ‘other’ group which consists of chlorophytes that could not be identified to the genus level (Figure 5c). The most common genera was *Bathycoccus*, the subclades A and B from clade VII of the prasinophytes (Lopes Dos Santos *et al*., 2017), which we refer to as Prasinophyte A and Prasinophyte B, and *Ostreococcus*. Prasinophyte A and B made up a large relative proportion at many of the sites (Figure 5c), consistent with other studies within the EEP region (Collado-Fabbri, Vaulot and Ulloa, 2011; de Vargas *et al*., 2015).

Spatial variation in chlorophyte communities was more apparent than shifts associated between years. Prasinophyte A, Prasinophyte B, and *Ostreococcus* were detected at all sites in 2015. Similarly in 2016, these groups were detected at all sites except at sites five, 22, and 24 which respectively lacked detection of either Prasinophyte B, *Ostreococcus*, or Prasinophyte A. Regardless of year, proportions of prasinophytes were generally highest in the east and slightly decreased westward (Figure 5c). *Ostreococcus* are cosmopolitan in protistan communities of the Peruvian coastal upwelling, and thus thrive under upwelling conditions, which could explain a higher detection at sites associated with the EUC (Collado-Fabbri, Vaulot and Ulloa, 2011; Rii *et al*., 2016). Moreover, total NO_3_^−^ uptake rates and picoeukaryote cell abundance (via flow cytometry) best explained changes in chlorophyte communities (Spearman’s rho = 0.6826) (Table 1); the latter being expected given that many chlorophytes are small enough to be enumerated as picoeukaryotes. Chlorophyte communities showed the most spatial changes relative to the dinoflagellates and diatoms, such that genera spread along the vertical axis of the NMDS plot (Figure 6).

The diatoms consisted of 78 identified OTUs within 15 genera. Diatom communities were well represented with seven genera and an ‘other’ group that mainly comprised of those which were unidentifiable to the genus level (Figure 5d). Diatoms were not detected through 18S sequencing at sites five or 26 in 2016 but may have been present at low abundances. Patterns of primary productivity by small cells and *Prochlorococcus* cell abundance provided the best prediction of diatom community change (Spearman’s rho = 0.3671) (Table 1). Given the patchy spatial extent of diatoms detected, the ability to predict the changes in the diatom community from the oceanographic variables was lower relative to the other groups.

Two common diatom genera*, Corethron* and *Pseudo-nitzschia* had slight temporal trends and are hypothesized to be somewhat regulated by the EUC upwelling conditions (Torres and Tapia, 2000; McCulloch, 2011). During 2015, *Pseudo-nitzschia* was present at the western sites (sites 3-5) and site 18. Similarly, during the 2006/07 El Niño, *Pseudo-nitzschia* was a dominant phytoplankton species north and west of Isabela Island (McCulloch, 2011). *Pseudo-nitzschia* was however still highly detected in 2016 at site 22. In prior neutral periods, this diatom genus has been observed in patchy distributions (Torres and Tapia, 2000; Tapia and Naranjo, 2012). *Corethron* followed opposite trends, such that it represented a higher proportion at sites 1, 3, and 4 in the neutral period while in 2015 it was only detected at site 4. Similarly, a 10-fold decrease in *Corethron* was measured during the 2006/07 El Niño relative to its highest abundance during cooler, neutral periods (McCulloch, 2011). *Pseudo-nitzschia* species are detected in the diatom community even in low nutrient conditions, likely due to physiological advantages, while *Corethron* are present in greater proportions at sites 1-4 in 2016, likely benefitting from the nutrient-rich neutral period. Two other ubiquitous diatom genera*, Chaetoceros* and *Thalassiosira,* were omnipresent in the GMR and have been previously identified during various seasons and stages of ENSO (Torres and Tapia, 1998; McCulloch, 2011; Tapia and Naranjo, 2012; Naranjo and Tapia, 2015). *Chaetoceros* can form resting spores anticipatory of upwelling relaxation (Pitcher *et al*., 1991), fair well under horizontal advection (Tilstone *et al*., 2015), and germinate rapidly (Smayda, 2000), which may explain their higher proportions at western sites in both 2015 and 2016 (Figure 5d).

### Deep water mass properties influence protist communities

The protistan community varied most between the El Niño and neutral periods, but also had some spatial patterns, such that western and eastern sites formed groupings (Figure 6). From the BIO-ENV analysis, this variation in protistan community composition is best explained (Spearman’s rho = 0.6239) collectively by patterns of total NO_3_^−^ uptake, picoeukaryote and *Prochlorococcus* cell abundances, and the density of subthermocline layer water masses (Table 1). Interestingly, *Synechococcus* abundance and density of the subthermocline layer had the highest BIO-ENV model predictability of any two variables combined (rho = 0.5664) (Table 1). Moreover, regressions between select individual variables and community dissimilarities showed that *Synechococcus* had the highest correlation with community change (R^2^ = 0.7271), due to its significant correlation with dinoflagellate and chlorophyte community subgroups (Supplementary Table 4). Dinoflagellates, the subgroup of the protistan community which varied the most, also had the strongest correlation with the density of the subthermocline layer (R^2^=0.73) (Supplementary Table 4).

In the broader EEP, phytoplankton biomass has been linked to changes in deep water mass conditions below the thermocline, such that decreases in Chl *a* biomass due to El Niño events can be detected before SST anomalies (Park, Dunne and Stock, 2018). Thermocline depth has also previously been identified as important for predicting Chl *a* concentrations in the GMR (Palacios, 2002; Sweet *et al*., 2007). In this study, the lack of strong correlation between the protistan communities with mixed layer properties is likely due to protists, particularly dinoflagellates, responding to deep water mass shifts before they are detected in the mixed layer. The EUC, a deep nutrient-rich current that slows during El Niño, not only upwells on the western side of the Galápagos platform, but flows horizontally around Isabela Island and continues eastward, providing deep water mass sources from the north and south to the archipelago (Jakoboski *et al*., 2020). The EUC likely influenced the observed spatio-temporal changes in protistan communities over the 2015/16 El Niño. These observations also provide support that protistan communities change in conjunction with cyanobacteria populations, and that the density of the subthermocline layer is a critical environmental indicator of that shift in community composition, both of which are attributes of the broader EEP (Masotti *et al*., 2010; Park, Dunne and Stock, 2018).

## Conclusions

Changes in water density profiles coupled with the nutrient regime suggest an observed weakening of the EUC, selecting for different phytoplankton size classes over the 2015/16 ENSO. In 2015, waters were less dense and had lower Si(OH)_4_:NO_3_^−^ ratios relative to the cooler, 2016 neutral period. Increased nutrient availability in 2016 likely led to increases in nitrate utilization by large cells such as diatoms and dinoflagellates. Despite appreciably lower primary productivity at western sites in 2015 compared to 2016, overall primary productivity of phytoplankton communities did not significantly differ across the entire archipelago due to local hotspots during El Niño (e.g., sites 14, 16) and small phytoplankton having higher primary productivity in 2015 at many stations, offsetting the higher NO_3_^−^ uptake in both small and large phytoplankton during 2016.

Protistan communities varied distinctly in the GMR during the 2015/16 ENSO. Chlorophytes were detected in high abundance in both years, varied spatially, and correlated with NO_3_^−^ uptake, picoeukaryote abundance, and *Synechococcus*. Diatoms had a patchy spatial extent making causes for changes difficult to discern, yet primary productivity by small cells and *Prochlorococcus* abundances were significant correlates. The largest difference between protistan communities however, was in the dinoflagellate group, such that *Syndiniales* were highly detected in 2015 while *Gyrodinium* were dominant in 2016. Dinoflagellates also happened to correlate strongly with primary productivity of small phytoplankton, picoeukaryote abundance and the density of the subthermocline layer.

The strongest correlation between the oceanographic variables and the entire protistan community composition over the 2015/16 ENSO was the density of the subthermocline layer – a proxy for shifts in deep water masses. These findings indicate that the water mass sources are an important factor in influencing protistan seed populations in the mixed layer whereas fluctuations in the short-term oceanographic conditions may have a more profound influence on their overall abundance and physiological status. Our observations provide motivation to continue to understand the effects of El Niño events on the microbial food-webs in the Galápagos Archipelago and the surrounding EEP region; specifically, to identify how changes in productivity and protistan community composition, as a function of altered ocean circulation, will influence higher marine trophic levels.

## Supplemental Information Additional Methods

### Phytoplankton biomass

Chl *a*, a proxy for phytoplankton biomass, was collected in triplicate by gravity filtering 400 ml of seawater through Isopore 5 µm polycarbonate filters (47 mm) to obtain the large size-fraction (> 5 µm). The filtrate was then filtered onto a Whatman GF/F filter (25mm) using an in-line vacuum (≤ 100 mmHg) to obtain the small size-fraction (≤ 5 µm). The filters were extracted in 6 ml of 90% acetone and incubated in the dark at −20 °C for 24 hours. Raw fluorescence values of the Chl *a* extracts were measured on a Turner Designs 10-AU fluorometer according to the methods of Brand et al. (1981). Dissolved inorganic nutrients (nitrate + nitrite, phosphate and silicic acid) were measured by filtering 30 ml of water through a 0.2 µm filter, using acid-washed syringes into a polypropylene Falcon^TM^ tube. Dissolved nutrient concentrations were analyzed using a OI Analytical Flow Solutions IV auto analyzer by Wetland Biogeochemistry Analytical Services at Louisiana State University.

Picophytoplankton counts, specifically *Prochlorococcus*, *Synechococcus*, and picoeukaryote populations were quantified using flow cytometry. Two ml whole seawater samples were collected at each depth and preserved using a 10% preservative cocktail consisting of 40% phosphate-buffered saline solution (PBS), 10% formalin, and 0.5% glutaraldehyde (Marie, Simon and Vaulot, 2005). Following 15 minutes on ice to allow the preservative to permeate membranes, samples were frozen at −20 °C until analysis onshore. *Prochlorococcus*, *Synechococcus*, and photosynthetic picoeukaryotes in seawater samples were enumerated using a BD FACSCalibur Flow Cytometer and populations characterized as previously described (Johnson *et al*., 2010). Briefly, cells were excited with a 488 nm laser (15 mW Ar) and inelastic forward (<15°) scatter, inelastic side (90°) scatter (SSC), green (530 ± 30 nm) fluorescence, orange fluorescence (585 ± 42 nm), and red fluorescence (> 670 nm) emissions were measured. Population mean properties (scatter and fluorescence) were normalized to 1.0 or 2.0 µm yellow green polystyrene beads (Polysciences, Warrington, PA).

### Particulate nutrient concentrations and biological uptake rates

Particulate nutrients, DIC and NO_3_^−^ uptake rates were sampled to assess water quality and phytoplankton productivity. Triplicate acid-washed polycarbonate bottles (618 ml) were filled with seawater from each of the four light depths and incubated on deck in tanks for 24 hr, beginning between 6:00 - 8:00am to capture photosynthesis and respiration cycles congruently across sites. The tanks were flushed with surface seawater via a flow-through system and covered with screening to mimic the incident irradiance depths at which the water samples were collected. Tracer isotope additions of ≤ 10% of ambient nutrient concentrations were used to measure uptake rates following methods described in Slawyk et al. (1977) (Slawyk, Collos and Auclair, 1977). For isotope additions in the field, ambient nitrate and bicarbonate were assumed to be 5 µM and 1200 µM, respectively. Nitrite concentration was assumed to be < 5% of ambient N, therefore N uptake rates were assumed to be that of nitrate. Nitrate uptake was measured by adding 0.5 µM ^15^N-labelled NO_3_^−^ to each bottle prior to incubation (Dugdale and Goering, 1967). Dissolved inorganic carbon (DIC) uptake rates were measured by adding 120 µM ^13^C-labeled HCO_3_ to each bottle prior to incubation (Hama *et al*., 1983). After incubation, the bottle contents were filtered to capture the plankton community at 24 hr of exposure to the trace isotopes. The large size-fraction (> 5 µm) was filtered onto a 5 µm polycarbonate filter (47 mm) and the remaining filtrate was filtered onto a pre-combusted (450 °C for 5 hours) GF/F (25 mm) to obtain the small size-fraction (≤ 5 µm). The particles trapped on the 5 µm polycarbonate filters were rinsed with particle-free (0.2 µm filtered) seawater onto a separate pre-combusted GF/F. The filters were dried for 24 to 48 hours in a combustion oven at 60 °C, wrapped in tin foil squares (30x30mm, Elemental Analysis D1067), pelletized, and stored in a desiccator. Filters were sent to the Stable Isotope Facility at University of California Davis for mass spectrometry analysis. Measurements of particulate nitrogen (PN) and particulate carbon (PC) were obtained simultaneously with uptake rates of nitrate and dissolved inorganic carbon (DIC). Dissolved nitrate concentrations, PN, PC and ^15^N and ^13^C atom percentages were used to calculate volumetric NO_3_^−^ uptake and DIC uptake rates of the different size fractions as according to Dugdale and Goering (1967) (Dugdale and Goering, 1967). Samples were not acidified to remove particulate inorganic carbon. NO_3_^−^ uptake rates were not corrected for possible losses of ^15^N in the form of dissolved organic nitrogen (Bronk, Glibert and Ward, 1994); therefore, the reported values are considered conservative estimates or net uptake. Chl *a*, particulate nutrients and nutrient uptake rates, as well as dissolved nutrient concentrations are displayed in Supplementary Table 3.

### Sequence library preparation for 18S amplicon sequencing

For protist taxonomic identification and proportions, four liters of seawater from 50% incident irradiance at the 18S sites (Figure 1a) was filtered using an in-line vacuum (< 100 mmHg) through 0.45 µm NES membrane filters (Pall, 47 mm). DNA was extracted from filters using the Qiagen DNeasy Plant Mini Kit (Qiagen, Germantown, MD, USA) and manufacturer provided protocol. DNA concentrations were quantified using the Quant-iT dsDNA High-Sensitivity Assay kit (Life Technologies, Carlsbad, CA, USA). Extracts were diluted either 1:10 or 1:100 so that the DNA concentrations were between 20-50 ng/µl.

The V4 hypervariable region of the 18S rRNA gene (600 bp) was targeted and amplified using a two-step PCR method described in Quigley et al., 2014. The forward linker primer (5’-<u>TCG TCG GCA GCG TC</u> + *A GAT GTG TAT AAG AGA CAG* + NNNN + **CCAGCASCYGCGGTAATTCC** -3’), and the reverse linker primer (5’-<u>GTC TCG TGG GCT</u> CGG + *AGA TGT GTA TAA GAG ACAG* + NNNN + **ACTTTCGTTCTTGAT** -3’) were used in the first PCR and barcodes were attached during the second PCR. The underlined nucleotide bases are the linker sequences, the italicized bases are spacer sequences, the N’s are degenerative nucleotide bases (which were not used for this study), and the bold bases are the V4-18S eukaryotic target sequences. The reagents used for the first PCR included 15 µl Milli-Q water, 2.4 µl ExTaq buffer, 1 µl of forward linker primer, 1 µl of reverse linker primer, 1 µl of ExTaq dNTPs, and 0.125 µl of ExTaq enzymes (Takara Bio Inc., Katsastu, Japan). Five µl of diluted DNA extract was added to the reaction mixture. Samples were run in a thermocycler at 95 °C for 5 min, 30 cycles at 95 °C for 40 s, 59 °C for 2 min, and 72 °C for 1 min, followed by a third stage at 72 °C for 7 min. Products of the reaction were checked on a 1% agarose gel. Products were cleaned using the Qiagen Qiaquick PCR purification kit (Qiagen, Germantown, MD, USA) and manufacturer provided protocol. The reagents used for the second PCR included 9.5 µl Milli-Q water, 3 µl of barcoded forward primer, 3 µl of barcoded reverse primer, 2 µl ExTaq buffer, 0.5 µl of ExTaq dNTPs, and 0.1 µl of ExTaq enzyme. Two µl of PCR product from the first reaction, diluted to 10 ng/µl was added to the reaction mixture. Samples were run in the thermocycler at 95 °C for 5 min, 4-10 cycles at 95 °C for 40 s, 59 °C for 2 min, and 72 °C for 1 min, followed by a third stage at 72 °C for 7 min. Samples were checked on a 1% agarose gel every 2 cycles until faint bands were achieved. Products were excised from the gel and cleaned using the Qiagen Gel Extraction kit (Qiagen, Germantown, MD, USA) and manufacturer provided protocol. DNA concentrations of the products were quantified, and samples were pooled each at a concentration of 10 ng/µl. The pool was run in a single large gel lane on a 1% SYBR Green (Invitrogen, Carlsbad, CA, USA) stained gel. The amplicon library was excised from the gel and submitted for sequencing to the University of North Carolina at Chapel Hill High Throughput Sequencing Facility across two lanes of Illumina MiSeq (2×300 base pairs).

---

**Supplementary Figure 1.**
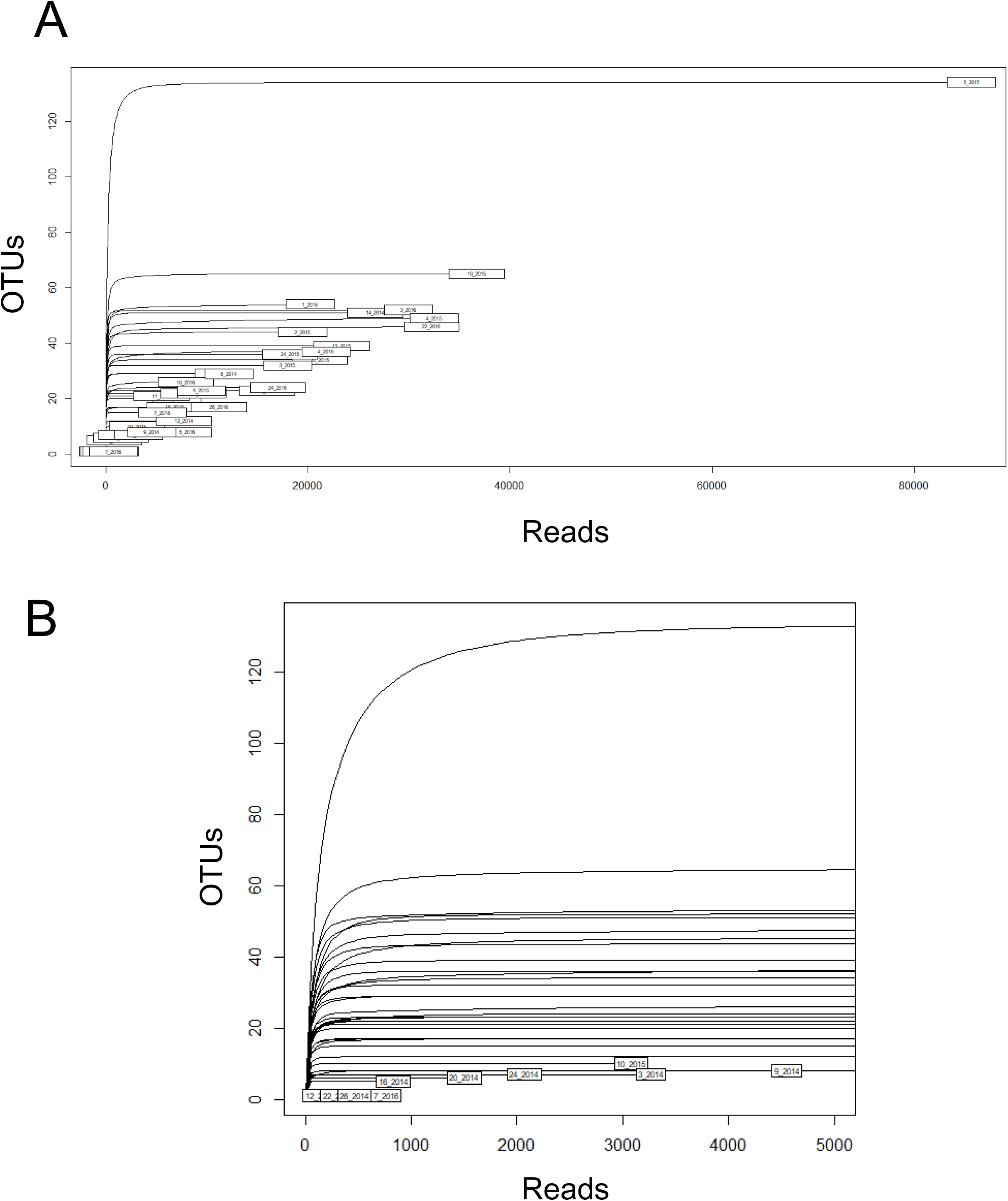
Rarefaction curves (technical replicates have been pooled). A) Rarefaction curves for 39 samples. B) Same as chart A, with x-axis set from 0 to 5000 reads. Samples were rarefied to 2066 reads. Note that OTU values on these plots reflect OTUs before custom taxonomy was assigned.

**Supplementary Figure 2.**
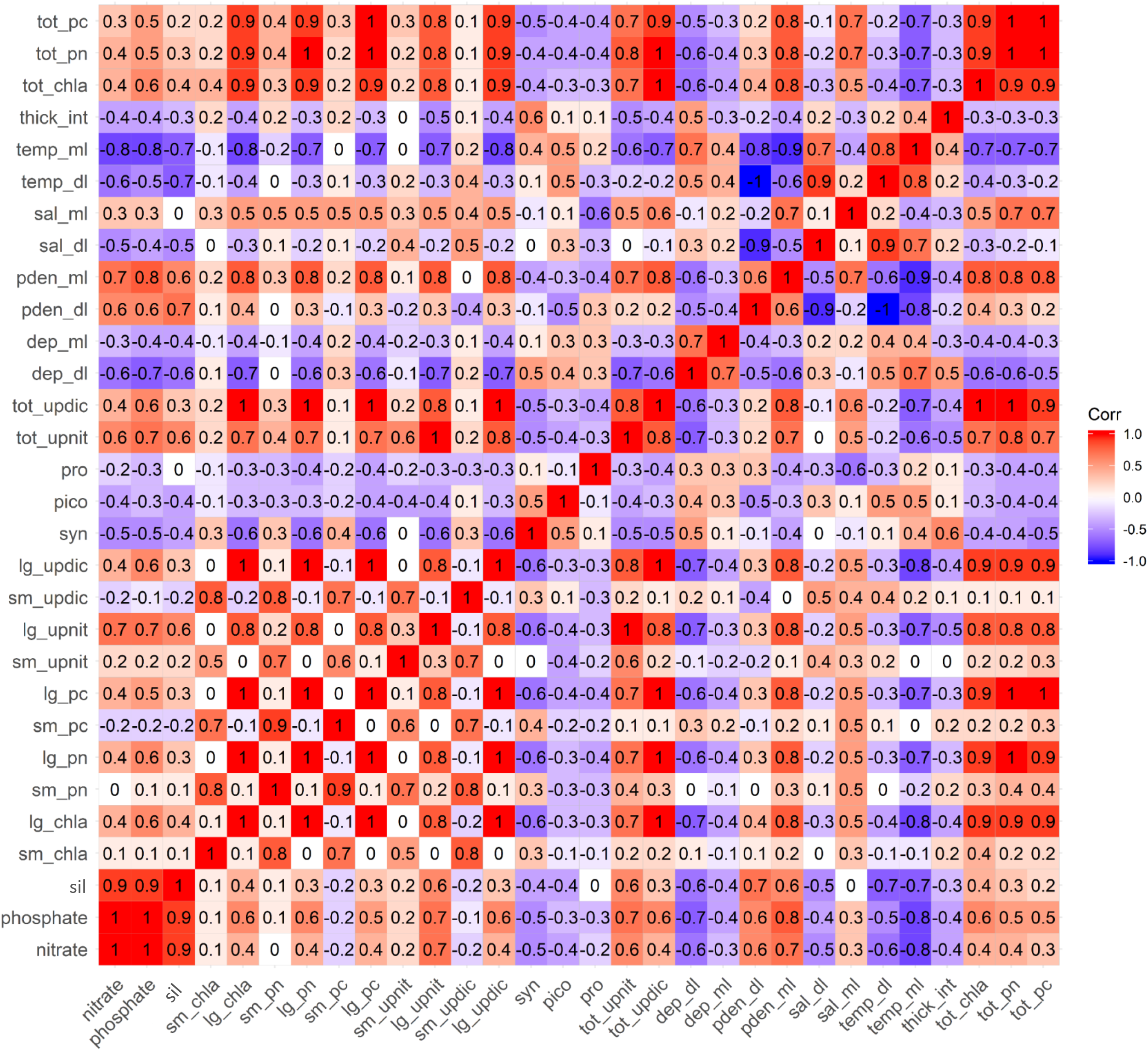
Correlation matrix of variables. Some variables are abbreviated such that: sil = silicic acid concentration; sm_chla = chlorophyll-a concentration (mg/L) of the small (<5 µm) phytoplankton size-fraction; lg_chla = chlorophyll-a concentration (mg/L) of the large (>5 µm) phytoplankton size-fraction; sm_pn = particulate nitrogen (<5 µm); lg_pn = particulate nitrogen (>5 µm); sm_pn = particulate carbon (<5 µm); lg_pn = particulate carbon (>5 µm); sm_upnit = nitrate uptake by the small (<5 µm) phytoplankton size-fraction; lg_upnit = nitrate uptake by the large (>5 µm) phytoplankton size-fraction; sm_updic = dissolved inorganic carbon uptake by the small (<5 µm) phytoplankton size-fraction; lg_updic = dissolved inorganic carbon uptake by the large (>5 µm) phytoplankton size-fraction; syn = *Synechococcus* flow-cytometry counts; pico = picoeukaryote flow-cytometry counts; pro = *Prochlorococcus* flow-cytometry counts; pden_dl = potential density of the subthermocline layer; tot_upnit = total nitrate uptake of both large and small phytoplankton size-fractions; tot_updic = total DIC uptake of both large and small phytoplankton size-fractions; dep_dl = depth of the subthermocline layer; dep_ml = depth of the mixed layer; pden_dl = density of the subthermocline layer; pden_ml = density of the mixed layer; sal_dl = salinity of the subthermocline layer; sal_ml = salinity of the mixed layer; temp_dl = temperature of the subthermocline layer; temp_ml = temperature of the mixed layer; thick_int = distance between the bottom of the mixed layer and top of the subthermocline layer (interfacial layer); tot_chla = total chlorophyll-a concentration (mg/L) of both large and small phytoplankton size-fractions; tot_pc = total particulate carbon of both large and small size-fractions; tot_pn = total particulate nitrogen of both large and small size-fractions.

**Supplementary Figure 3.**
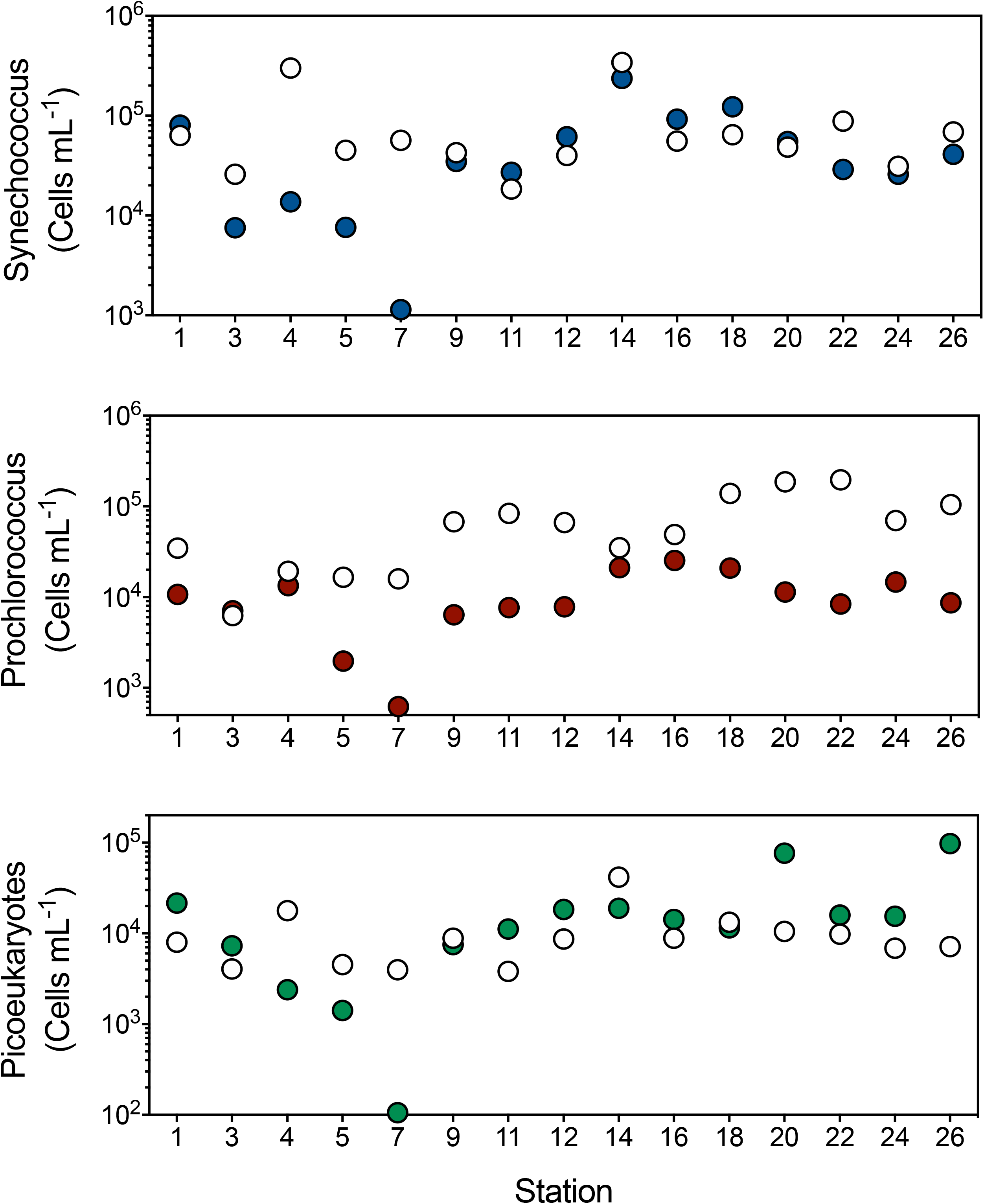
Flow cytometry cell counts at sampling sites in 2015 and 2016. White circles indicate cell densities in 2015, while colored circles indicate cell densities in 2016.

**Supplementary Figure 4.**
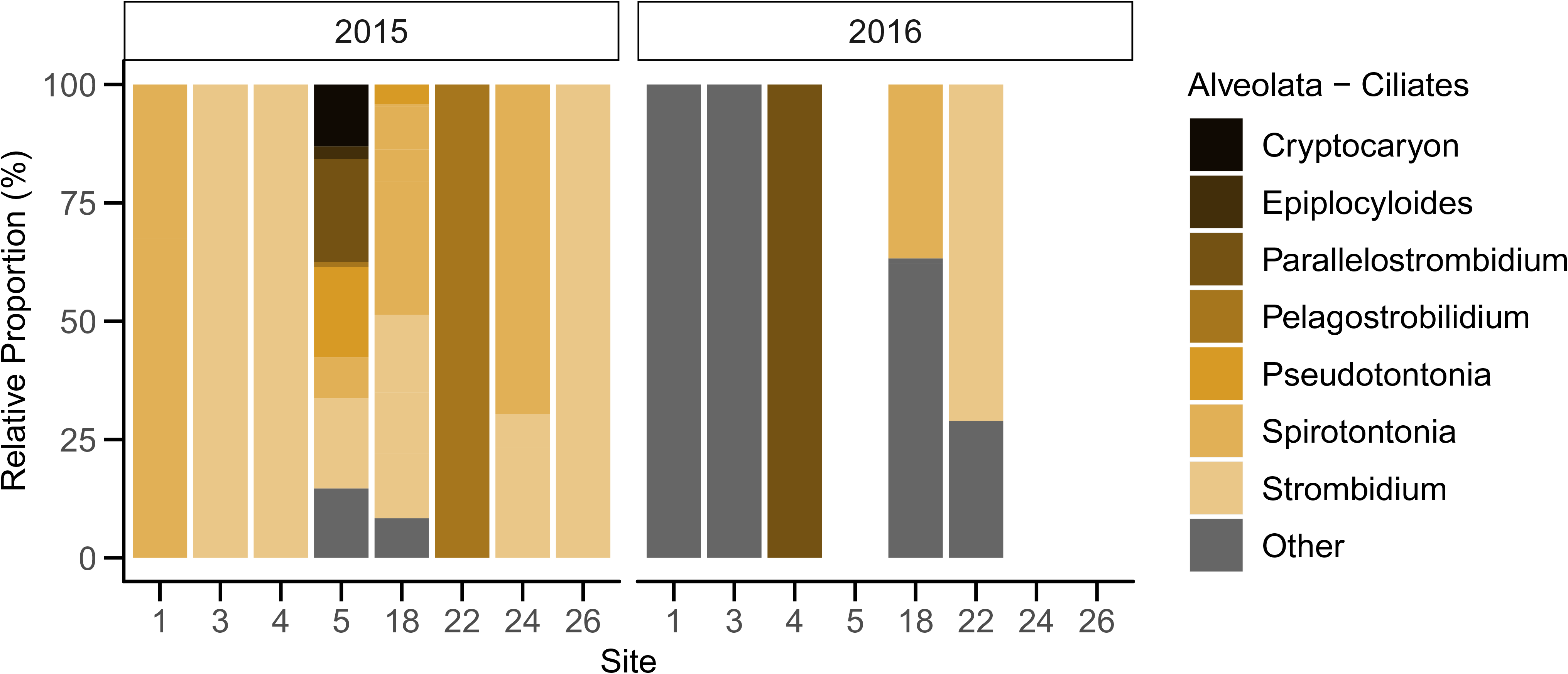
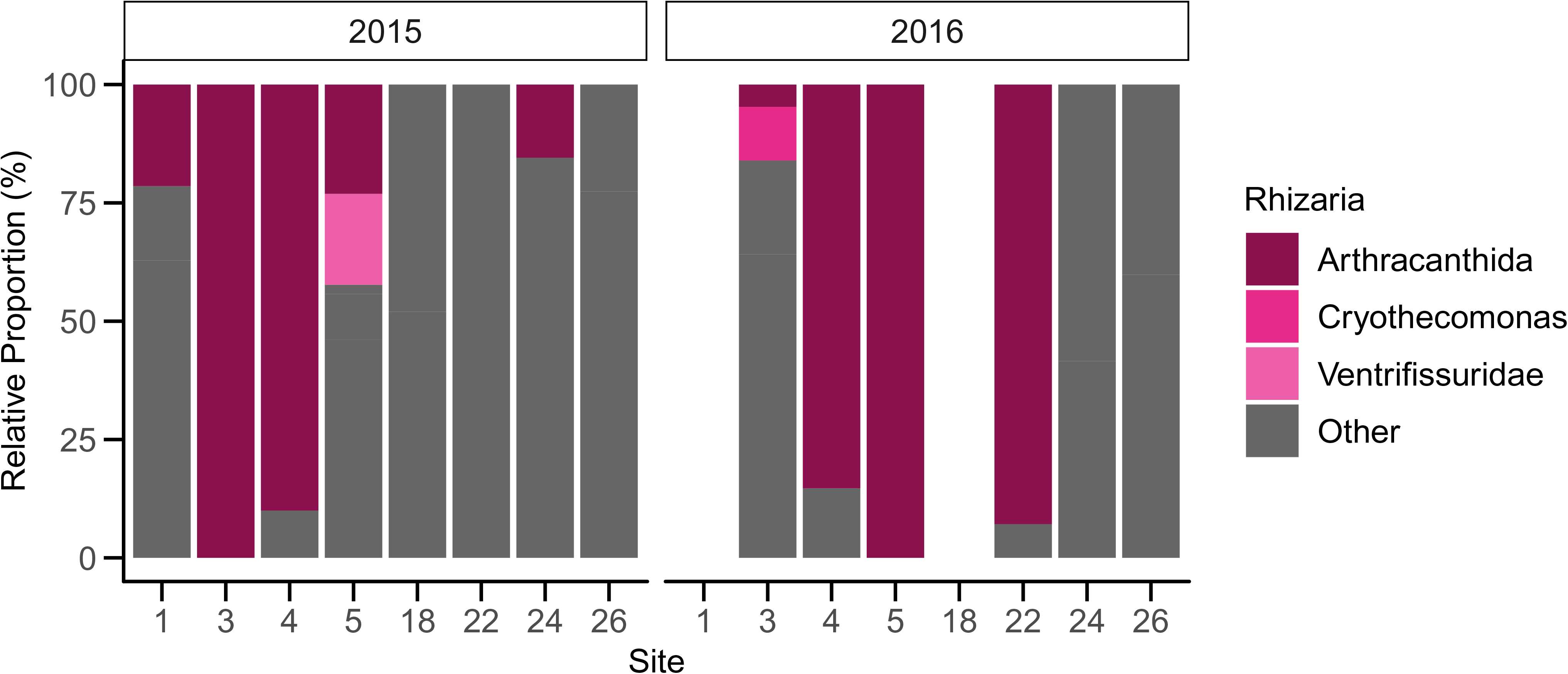
Ciliate and Rhizaria 18S rRNA gene community plots. A) Relative proportions of the Ciliate group, highlighting the top seven most abundant genera. B) Relative proportions of the Rhizaria group, highlighting the top three most abundant genera.

**Supplementary Table 1.** Table of samples used for 18S rRNA gene analysis of ‘18S’ sites in this study. Samples are derived from bolded samples in Appendix 1, some of which have been pooled from technical replicates to obtain the values below. The final number of reads, or library size, was obtained after blasting the assembled amplicons to the SILVA v. 123 reference database. The number of different OTUs present in each sample before and after rarefication is displayed. OTUs no. values reflect adjustments made to unknown OTUs. If unknown OTUs were present in the same Class they were classified as the same unknown grouping and reflected a single OTU.

| Site | Year | Reads |  |  |  |  | OTUs |  |  |
| --- | --- | --- | --- | --- | --- | --- | --- | --- | --- |
|  |  | input | filtered | denoised | merged | non-chimeric | final no. of reads | Raw | After Rarefy |
| 1 | 2015 | 60327 | 27917 | 27917 | 13158 | 10576 | 7044 | 14 | 14 |
| 1 | 2016 | 191084 | 152385 | 152385 | 130136 | 28722 | 20227 | 15 | 15 |
| 3 | 2015 | 176261 | 52398 | 52398 | 24918 | 19463 | 17994 | 17 | 17 |
| 3 | 2016 | 86983 | 71192 | 71192 | 56869 | 33540 | 29945 | 26 | 26 |
| 4 | 2015 | 149983 | 120461 | 120461 | 94927 | 43158 | 32498 | 19 | 19 |
| 4 | 2016 | 130178 | 76059 | 76059 | 65219 | 26087 | 21793 | 19 | 19 |
| 5 | 2015 | 845886 | 761534 | 761534 | 589629 | 145115 | 85659 | 56 | 53 |
| 5 | 2016 | 21114 | 17367 | 17367 | 14640 | 11223 | 8063 | 8 | 8 |
| 18 | 2015 | 188403 | 100918 | 100918 | 59478 | 45265 | 36694 | 26 | 24 |
| 18 | 2016 | 62233 | 45655 | 45655 | 37554 | 8677 | 7928 | 10 | 10 |
| 22 | 2015 | 93016 | 53299 | 53299 | 33250 | 23350 | 15958 | 12 | 12 |
| 22 | 2016 | 187324 | 171043 | 171043 | 116206 | 46941 | 32238 | 22 | 22 |
| 24 | 2015 | 100552 | 66927 | 66927 | 42605 | 26749 | 18215 | 20 | 20 |
| 24 | 2016 | 78095 | 71395 | 71395 | 52183 | 22001 | 17021 | 12 | 12 |
| 26 | 2015 | 51114 | 34406 | 34406 | 20306 | 17974 | 11148 | 12 | 12 |
| 26 | 2016 | 43342 | 37709 | 37709 | 27984 | 19119 | 11163 | 10 | 10 |

**Supplementary Table 2.**
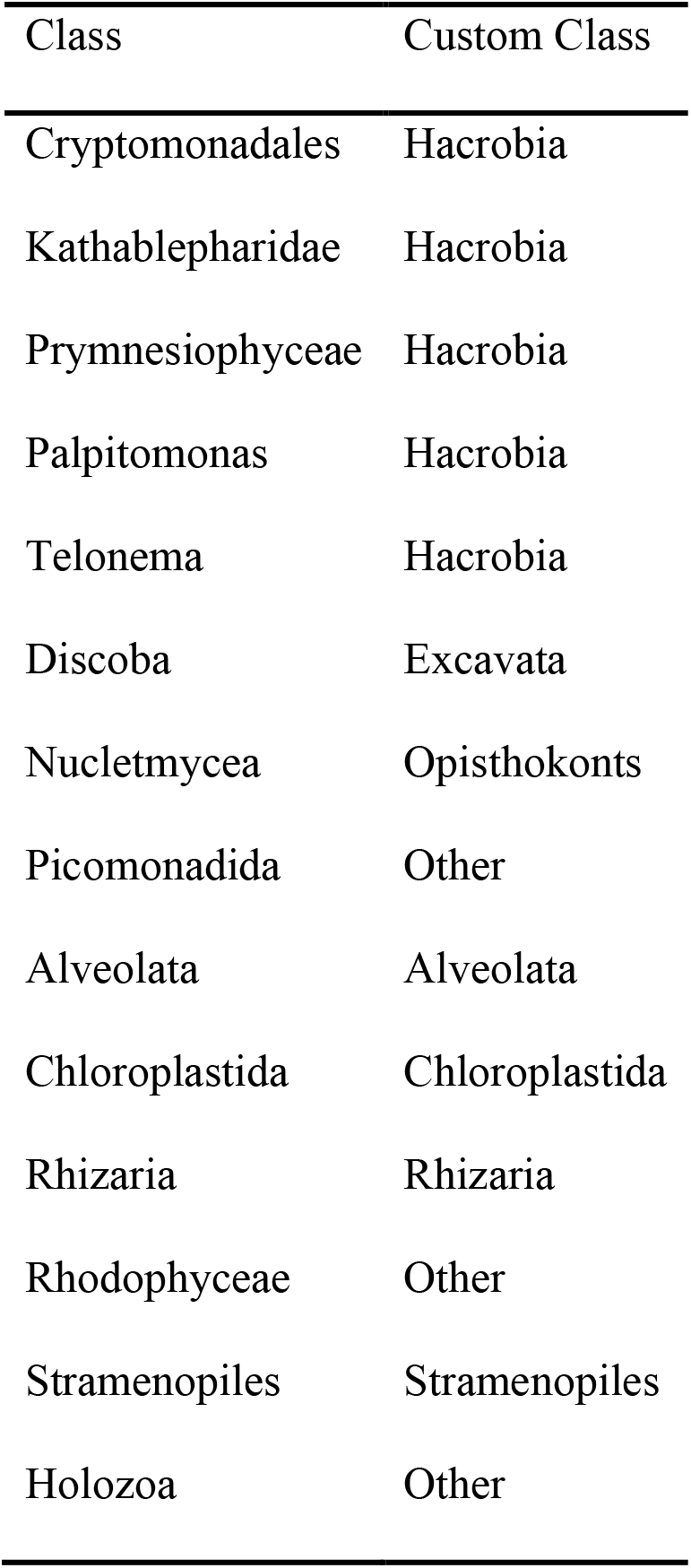
Custom taxonomy assigned to the Class taxonomic level (D_2) originating from the SILVA v. 123 database.

**Supplementary Table 3.** Summary statistics for dissolved and particulate nutrients, Chl *a*, primary productivity and nitrate uptake rates. See supplementary Figure 2 for abbreviated variable key.

| Variable | mean |  | median |  | independent 2-group Mann-Whitney U Test |  |
| --- | --- | --- | --- | --- | --- | --- |
|  | 2015 | 2016 | 2015 | 2016 | W | p-value |
| nitrate | 2.157467 | 6.758967 | 1.6995 | 6.193 | 29 | 0.000264 |
| silicic acid | 1.808267 | 6.081933 | 1.654 | 4.985 | 12 | 3.51E-06 |
| sm_chla | 0.334206 | 0.349624 | 0.224223 | 0.230617 | 93 | 0.4306 |
| lg_chla | 0.196312 | 0.422502 | 0.125355 | 0.200933 | 67 | 0.06128 |
| tot_chla | 0.528531 | 0.772019 | 0.407707 | 0.637963 | 72 | 0.09753 |
| sm_updic | 2.442711 | 1.962444 | 1.901882 | 1.436667 | 132 | 0.4363 |
| lg_updic | 1.280083 | 2.673333 | 0.462093 | 1.2 | 78 | 0.1607 |
| tot_updic | 3.707268 | 4.561556 | 2.677045 | 3.146667 | 89 | 0.3453 |
| sm_upnit | 198.8752 | 159.2167 | 81.37914 | 129.8633 | 108 | 0.8702 |
| lg_upnit | 169.0535 | 342.1778 | 51.91571 | 131.64 | 69 | 0.0742 |
| tot_upnit | 367.9203 | 494.6082 | 105.1708 | 329.3867 | 81 | 0.2017 |

| year | latitude | longitude | site | NO<sub>3</sub><sup>-</sup><br>(μM) | PO<sub>4</sub><sup>3-</sup><br>(μM) | Si(OH)<sub>4</sub><br>(μM) |
| --- | --- | --- | --- | --- | --- | --- |
| 2015 | -1.08327 | -91.0913 | 1 | 5.0165 | 0.523 | 2.4385 |
| 2015 | -0.46945 | -91.6154 | 3 | 5.6755 | 0.796 | 3.268 |
| 2015 | -0.44772 | -91.3761 | 4 | 1.074 | 0.408 | 3.1975 |
| 2015 | -0.28752 | -91.6564 | 5 | 1.6995 | 0.4345 | 1.017 |
| 2015 | -0.06255 | -91.5572 | 7 | 0.5475 | 0.5735 | 3.0915 |
| 2015 | 0.160683 | -91.3097 | 9 | 2.853 | 0.4845 | 2.0715 |
| 2015 | 0.548283 | -90.7919 | 11 | 0.193 | 0.3735 | 1.618 |
| 2015 | 0.286917 | -90.5176 | 12 | 2.6645 | 0.371 | 1.022 |
| 2015 | 0.306183 | -89.9517 | 14 | 0.7 | 0.294 | 0.613 |
| 2015 | -0.22822 | -90.8716 | 16 | 3.851 | 0.478 | 1.654 |
| 2015 | -0.36595 | -90.5421 | 18 | 2.3585 | 0.5485 | 1.328 |
| 2015 | -0.55573 | -90.1461 | 20 | 1.48 | 0.508 | 2.31 |
| 2015 | -1.20072 | -90.4423 | 22 | 2.637 | 0.5735 | 2.249 |
| 2015 | -1.33905 | -89.7414 | 24 | 1.478 | 0.445 | 0.9935 |
| 2015 | -0.68873 | -89.2281 | 26 | 0.134 | 0.383 | 0.2525 |
| 2016 | -1.08327 | -91.0913 | 1 | 3.232 | 0.524 | 2.902 |
| 2016 | -0.46945 | -91.6154 | 3 | 17.304 | 1.4265 | 14.0895 |
| 2016 | -0.44772 | -91.3761 | 4 | 9.2245 | 0.901 | 7.3305 |
| 2016 | -0.28752 | -91.6564 | 5 | 7.7885 | 0.789 | 4.985 |
| 2016 | -0.06255 | -91.5572 | 7 | 2.908 | 0.4855 | 1.7345 |
| 2016 | 0.160683 | -91.3097 | 9 | 5.603 | 0.6195 | 4.098 |
| 2016 | 0.548283 | -90.7919 | 11 | 8.3315 | 0.7935 | 6.599 |
| 2016 | 0.286917 | -90.5176 | 12 | 3.7985 | 0.669 | 3.126 |
| 2016 | 0.306183 | -89.9517 | 14 | 2.344 | 0.6065 | 4.193 |
| 2016 | -0.22822 | -90.8716 | 16 | 8.335 | 0.828 | 8.4035 |
| 2016 | -0.36595 | -90.5421 | 18 | 6.3325 | 0.7765 | 6.375 |
| 2016 | -0.55573 | -90.1461 | 20 | 4.0875 | 0.5285 | 4.876 |
| 2016 | -1.20072 | -90.4423 | 22 | 6.193 | 0.6425 | 6.594 |
| 2016 | -1.33905 | -89.7414 | 24 | 14.535 | 1.203 | 11.628 |
| 2016 | -0.68873 | -89.2281 | 26 | 1.3675 | 0.4045 | 4.295 |

|  |  | mean | mean | mean | standard<br>deviation | standard<br>deviation | standard<br>deviation |
| --- | --- | --- | --- | --- | --- | --- | --- |
| year | site | sm_pn<br>(nmol/L) | lg_pn<br>(nmol/L) | tot_pn<br>(nmol/L) | sm_pn | lg_pn | tot_pn |
| 2015 | 1 | 1392.873 | 524.3925 | 1917.266 | 197.2441 | 185.0112 | 193.3588 |
| 2015 | 3 | 1263.166 | 1449.514 | 2712.68 | 266.6868 | 245.371 | 200.0526 |
| 2015 | 4 | 1709.851 | 597.4289 | 2307.28 | 191.5843 | 218.3932 | 26.80889 |
| 2015 | 5 | 1325.54 | 1069.004 | 2394.543 | 104.2701 | 31.07704 | 100.2418 |
| 2015 | 7 | 906.2676 | 770.768 | 1677.036 | 512.5574 | 130.8078 | 389.7119 |
| 2015 | 9 | 885.3941 | 650.482 | 1535.876 | 73.24488 | 421.7521 | 414.527 |
| 2015 | 11 | 935.1361 | 340.5332 | 1275.669 | 136.0965 | 43.05484 | 130.1387 |
| 2015 | 12 | 695.9137 | 466.6674 | 1162.581 | 376.425 | 50.11469 | 425.5941 |
| 2015 | 14 | 2528.893 | 470.1441 | 2999.037 | 196.9775 | 125.9723 | 100.7986 |
| 2015 | 16 | 1799.034 | 1804.49 | 3603.525 | 339.6629 | 548.5873 | 408.6837 |
| 2015 | 18 | 1271.441 | 587.7022 | 1859.143 | 122.6524 | 240.593 | 159.2031 |
| 2015 | 20 | 976.7436 | 523.3313 | 1500.075 | 519.1229 | 459.3802 | 169.1233 |
| 2015 | 22 | 954.3289 | 745.4182 | 1699.747 | 122.2689 | 430.922 | 500.2987 |
| 2015 | 24 | 927.9323 | 781.1187 | 1709.051 | 553.4452 | 518.2618 | 84.22699 |
| 2015 | 26 | 914.6053 | 558.8385 | 1473.444 | 85.72798 | 50.27337 | 98.89218 |
| 2016 | 1 | 1373.89 | 783.57 | 2157.46 | 33.51172 | 165.3045 | 147.6448 |
| 2016 | 3 | 1374.937 | 2229.037 | 3603.973 | 178.4089 | 88.74901 | 146.9314 |
| 2016 | 4 | 1549.96 | 3677.58 | 5227.54 | NA | NA | NA |
| 2016 | 5 | 1036.717 | 2463.55 | 3500.267 | 97.76806 | 288.4142 | 222.2838 |
| 2016 | 7 | 1860.9 | 2285.52 | 4146.42 | 391.312 | 223.41 | 301.7416 |
| 2016 | 9 | 982.0933 | 2017.673 | 2999.767 | 206.0789 | 158.0138 | 68.70205 |
| 2016 | 11 | 1131.64 | 848.8333 | 1226.047 | NA | 55.91428 | 663.4499 |
| 2016 | 12 | 1068.987 | 949.3067 | 2018.293 | 52.88796 | 306.0843 | 355.8799 |
| 2016 | 14 | 2511.533 | 726.62 | 3238.153 | 217.9735 | 89.13549 | 142.8141 |
| 2016 | 16 | 1990.683 | 311.74 | 2302.423 | 98.49966 | 49.06905 | 146.9604 |
| 2016 | 18 | 1794.35 | 463.16 | 2257.51 | 41.18031 | 166.6094 | 171.6449 |
| 2016 | 20 | 1052.63 | 603.27 | 1655.9 | 79.62618 | 334.6954 | 402.3871 |
| 2016 | 22 | 1051.527 | 589.79 | 1641.317 | 28.63359 | 19.17819 | 31.44936 |
| 2016 | 24 | 894.7867 | 465.64 | 1360.427 | 78.99461 | 63.19213 | 90.50729 |
| 2016 | 26 | 859.48 | 262.4467 | 1121.927 | 116.3601 | 1.403436 | 117.4547 |
| | site | sm_pc<br>($\mu\text{mol/L}$) | lg_pc<br>($\mu\text{mol/L}$) | tot_pc<br>($\mu\text{mol/L}$) | sm_pc | lg_pc | tot_pc |
| 2015 | 1 | 9.012001 | 2.986874 | 11.99888 | 3.130695 | 0.806295 | 2.392851 |
| 2015 | 3 | 7.221374 | 7.986993 | 15.20837 | 1.736978 | 1.577957 | 1.762298 |
| 2015 | 4 | 9.094588 | 4.202203 | 13.29679 | 0.731026 | 3.01544 | 2.284415 |
| 2015 | 5 | 7.280883 | 7.194916 | 14.4758 | 0.70243 | 0.4825 | 0.681973 |
| 2015 | 7 | 5.452462 | 4.21298 | 9.665441 | 3.610194 | 1.064037 | 2.569136 |
| 2015 | 9 | 5.398573 | 4.390504 | 9.789077 | 0.642065 | 1.530488 | 1.842264 |
| 2015 | 11 | 5.435372 | 1.733863 | 7.169235 | 2.25321 | 0.375027 | 2.295055 |
| 2015 | 12 | 3.265884 | 2.339255 | 5.60514 | 1.690878 | 0.461255 | 1.942756 |
| 2015 | 14 | 18.40962 | 3.196466 | 21.60609 | 1.080984 | 0.941431 | 0.673006 |
| 2015 | 16 | 8.67628 | 11.36877 | 20.04505 | 0.630804 | 3.637747 | 4.039047 |
| 2015 | 18 | 7.515143 | 2.652905 | 10.16805 | 0.50914 | 2.048071 | 2.279657 |
| 2015 | 20 | 5.249063 | 2.658915 | 7.907978 | 2.420769 | 2.532813 | 0.782465 |
| 2015 | 22 | 5.715023 | 3.106268 | 8.821291 | 0.715471 | 2.178608 | 2.455808 |
| 2015 | 24 | 4.614055 | 3.04495 | 7.659005 | 2.775506 | 2.341621 | 0.481262 |
| 2015 | 26 | 4.994912 | 2.326362 | 7.321275 | 0.629223 | 0.459367 | 0.177925 |
| 2016 | 1 | 7.948667 | 4.270667 | 12.21933 | 0.276685 | 0.290676 | 0.392538 |
| 2016 | 3 | 6.105 | 12.52467 | 18.62967 | 0.396849 | 0.488686 | 0.113271 |
| 2016 | 4 | 8.564 | 19.64 | 28.204 | NA | NA | NA |
| 2016 | 5 | 5.837333 | 14.31267 | 20.15 | 0.448536 | 2.132606 | 1.929112 |
| 2016 | 7 | 9.226 | 14.94533 | 24.17133 | 1.555803 | 2.119677 | 1.038851 |
| 2016 | 9 | 5.136667 | 10.45533 | 15.592 | 0.499829 | 0.187961 | 0.332608 |
| 2016 | 11 | 6.67 | 4.841 | 7.064333 | NA | 0.732885 | 3.80346 |
| 2016 | 12 | 6.918667 | 4.926667 | 11.84533 | 1.163151 | 0.882565 | 2.026655 |
| 2016 | 14 | 14.16233 | 4.550667 | 18.713 | 0.940403 | 1.000958 | 1.253807 |
| 2016 | 16 | 10.67267 | 1.667667 | 12.34033 | 0.387807 | 0.245875 | 0.525508 |
| 2016 | 18 | 9.140667 | 2.351 | 11.49167 | 0.204671 | 0.293714 | 0.280293 |
| 2016 | 20 | 5.861333 | 3.766333 | 9.627667 | 0.397649 | 2.339338 | 2.421094 |
| 2016 | 22 | 5.673333 | 3.489667 | 9.163 | 0.848373 | 0.502693 | 0.759947 |
| 2016 | 24 | 4.616333 | 2.397 | 7.013333 | 0.553079 | 0.310598 | 0.264434 |
| 2016 | 26 | 4.930333 | 1.583333 | 6.513667 | 0.684293 | 0.137758 | 0.818881 |
| year | site | sm_chla<br>(µg/L) | lg_chla<br>(µg/L) | tot_chla<br>(µg/L) | sm_chla | lg_chla | tot_chla |
| 2015 | 1 | 0.271717 | 0.135991 | 0.407707 | 0.027718 | 0.016219 | 0.026048 |
| 2015 | 3 | 0.236097 | 0.308935 | 0.545032 | 0.030888 | 0.117746 | 0.101418 |
| 2015 | 4 | 0.62061 | 0.125355 | 0.745965 | 0.158334 | 0.00535 | 0.163629 |
| 2015 | 5 | 0.264666 | 0.353442 | 0.618107 | 0.018209 | 0.022634 | 0.039772 |
| 2015 | 7 | 0.211 | 0.256 | 0.467 | 0.016371 | 0.015524 | 0.025239 |
| 2015 | 9 | 0.095215 | 0.070555 | 0.162551 | 0.008719 | 0.00567 | 0.007266 |
| 2015 | 11 | 0.074893 | 0.054252 | 0.129145 | 0.057838 | 0.005434 | 0.053563 |
| 2015 | 12 | 0.133411 | 0.09036 | 0.226754 | 0.000736 | 0.007499 | 0.006948 |
| 2015 | 14 | 1.32205 | 0.159377 | 1.47686 | 0.009687 | 0.014649 | 0.027125 |
| 2015 | 16 | 0.48224 | 0.885477 | 1.344655 | 0.009687 | 0.040417 | 0.000969 |
| 2015 | 18 | 0.555307 | 0.175817 | 0.731123 | 0.168015 | 0.01702 | 0.152319 |
| 2015 | 20 | 0.212504 | 0.069687 | 0.282192 | 0.021862 | 0.002093 | 0.021853 |
| 2015 | 22 | 0.17125 | 0.053247 | 0.222557 | 0.021312 | 0.006255 | 0.028771 |
| 2015 | 24 | 0.224223 | 0.087908 | 0.312132 | 0.071481 | 0.010183 | 0.079966 |
| 2015 | 26 | 0.137913 | 0.118277 | 0.25619 | 0.059816 | 0.009122 | 0.061939 |
| 2016 | 1 | 0.24249 | 0.119601 | 0.362091 | 0.014434 | 0.016421 | 0.030719 |
| 2016 | 3 | 0.310533 | 0.981377 | 1.29191 | 0.044315 | 0.147517 | 0.189035 |
| 2016 | 4 | 0.544347 | 0.91516 | 1.459507 | 0.389951 | 0.298657 | 0.16115 |
| 2016 | 5 | 0.184493 | 0.94119 | 1.124085 | 0.013306 | 0.156935 | 0.138529 |
| 2016 | 7 | 0.21783 | 1.429367 | 1.647197 | 0.03858 | 0.064744 | 0.101476 |
| 2016 | 9 | 0.184493 | 0.45347 | 0.637963 | 0.030828 | 0.007249 | 0.035205 |
| 2016 | 11 | 0.230617 | 0.287243 | 0.51786 | 0.021061 | 0.026577 | 0.023887 |
| 2016 | 12 | 0.202303 | 0.194997 | 0.3973 | 0.012729 | 0.019963 | 0.032304 |
| 2016 | 14 | 0.951237 | 0.27263 | 1.223867 | 0.092719 | 0.07475 | 0.12273 |
| 2016 | 16 | 0.649837 | 0.105901 | 0.755738 | 0.040925 | 0.009882 | 0.042926 |
| 2016 | 18 | 0.621067 | 0.089918 | 0.710984 | 0.037577 | 0.003839 | 0.033855 |
| 2016 | 20 | 0.216917 | 0.109874 | 0.326791 | 0.031161 | 0.013543 | 0.033058 |
| 2016 | 22 | 0.17262 | 0.200933 | 0.373553 | 0.041802 | 0.009623 | 0.050973 |
| 2016 | 24 | 0.336107 | 0.16166 | 0.497767 | 0.030888 | 0.014303 | 0.031905 |
| 2016 | 26 | 0.17947 | 0.074208 | 0.253678 | 0.0214 | 0.019151 | 0.03997 |
| year | site | sm_updic<br>($\mu\text{mol/L/day}$) | lg_updic<br>($\mu\text{mol/L/day}$) | tot_updic<br>($\mu\text{mol/L/day}$) | sm_updic | lg_updic | tot_updic |
| 2015 | 1 | 1.901882 | 0.462093 | 2.363976 | 0.173273 | 0.126476 | 0.140799 |
| 2015 | 3 | 2.531358 | 2.800502 | 5.33186 | 0.10659 | 0.490954 | 0.4881 |
| 2015 | 4 | 2.915005 | 0.498011 | 3.413016 | 0.41805 | 0.090995 | 0.327054 |
| 2015 | 5 | 2.013047 | 2.476068 | 4.489115 | 0.101204 | 0.158936 | 0.129727 |
| 2015 | 7 | 1.83 | 1.21 | 2.944496 | 0.06 | 0.07 | NA |
| 2015 | 9 | 1.22093 | 0.25 | 1.453445 | 0.11127 | 0.01 | 0.143663 |
| 2015 | 11 | 1.116283 | 0.166385 | 1.282669 | 0.209437 | 0.055016 | 0.196753 |
| 2015 | 12 | 0.77 | 0.271957 | 1.05045 | 0.1838 | 0.022101 | 0.161292 |
| 2015 | 14 | 9.586615 | 1.08495 | 10.67157 | 0.658968 | 0.295943 | 0.408357 |
| 2015 | 16 | 3.634229 | 8.14 | 11.70804 | 0.302435 | 1.393 | 1.787397 |
| 2015 | 18 | 2.735525 | 0.392058 | 3.127583 | 0.079292 | 0.293056 | 0.313838 |
| 2015 | 20 | 1.37 | 0.16 | 1.494396 | 0.0289 | 0.0643 | 0.020774 |
| 2015 | 22 | 1.353744 | 0.394033 | 1.747777 | 0.246938 | 0.339962 | 0.413958 |
| 2015 | 24 | 2.39 | 0.313642 | 2.677045 | 0.28 | 0.072618 | 0.358021 |
| 2015 | 26 | 1.272052 | 0.581541 | 1.853593 | 0.074504 | 0.037884 | 0.106918 |
| 2016 | 1 | 1.436667 | 0.553333 | 1.99 | 0.045092 | 0.095044 | 0.086603 |
| 2016 | 3 | 1.463333 | 6.116667 | 7.58 | 0.153731 | 0.332466 | 0.362905 |
| 2016 | 4 | 2.49 | 11.64 | 14.13 | NA | NA | NA |
| 2016 | 5 | 0.883333 | 5.48 | 6.363333 | 0.102632 | 0.167033 | 0.183394 |
| 2016 | 7 | 1.066667 | 5.176667 | 6.243333 | 0.106927 | 0.202073 | 0.106927 |
| 2016 | 9 | 1.156667 | 4.52 | 5.676667 | 0.202567 | 0.420357 | 0.228108 |
| 2016 | 11 | 1.67 | 1.383333 | 1.94 | NA | 0.120554 | 0.86885 |
| 2016 | 12 | 1.583333 | 1.2 | 2.783333 | 0.150444 | 0.147309 | 0.282902 |
| 2016 | 14 | 6.636667 | 1.616667 | 8.253333 | 0.768722 | 0.245832 | 0.645316 |
| 2016 | 16 | 3.406667 | 0.24 | 3.646667 | 0.106927 | 0.036056 | 0.12741 |
| 2016 | 18 | 2.75 | 0.396667 | 3.146667 | 0.121244 | 0.075056 | 0.058595 |
| 2016 | 20 | 1.406667 | 0.39 | 1.796667 | 0.068069 | 0.07 | 0.10504 |
| 2016 | 22 | 1.42 | 0.8 | 2.22 | 0.117898 | 0.079373 | 0.144222 |
| 2016 | 24 | 1.243333 | 0.473333 | 1.716667 | 0.140119 | 0.083865 | 0.105987 |
| 2016 | 26 | 0.823333 | 0.113333 | 0.936667 | 0.083267 | 0.015275 | 0.097125 |
| year | site | sm_upnit<br>(nmol/L/day) | lg_upnit<br>(nmol/L/day) | tot_upnit<br>(nmol/L/day) | sm_upnit | lg_upnit | tot_upnit |
| 2015 | 1 | 126.537 | 65.85543 | 192.3924 | 7.746767 | 14.21564 | 20.66745 |
| 2015 | 3 | 360.649 | 406.502 | 767.151 | 76.23357 | 163.9518 | 191.7983 |
| 2015 | 4 | 481.5605 | 117.6843 | 599.2448 | 57.97254 | 5.541328 | 52.43121 |
| 2015 | 5 | 255.933 | 387.2819 | 643.2149 | 22.77127 | 36.08576 | 40.91201 |
| 2015 | 7 | 145.48 | 164.8 | 330.6916 | 25.8 | 3.07 | NA |
| 2015 | 9 | 68.6046 | 27.83 | 99.93725 | 11.23629 | 0.43 | 13.81445 |
| 2015 | 11 | 50.73533 | 16.99245 | 67.72778 | 3.744021 | 2.951693 | 3.147611 |
| 2015 | 12 | 33.72984 | 42.96051 | 76.69035 | 17.56143 | 1.327783 | 16.71017 |
| 2015 | 14 | 700.0781 | 89.2328 | 789.3109 | 50.56118 | 27.78318 | 31.92963 |
| 2015 | 16 | 442.6527 | 1083.56 | 1501.049 | 52.54894 | 171.5795 | 213.1177 |
| 2015 | 18 | 81.37914 | 23.79163 | 105.1708 | 5.995178 | 10.4649 | 11.39536 |
| 2015 | 20 | 52.09 | 11.99 | 64.87759 | 4.728152 | 2.171333 | 5.795378 |
| 2015 | 22 | 62.03894 | 23.87998 | 85.91892 | 16.68338 | 12.30874 | 22.79962 |
| 2015 | 24 | 71.54 | 21.52602 | 93.39189 | 18 | 6.112541 | 26.60459 |
| 2015 | 26 | 50.11906 | 51.91571 | 102.0348 | 9.799634 | 5.009725 | 14.40122 |
| 2016 | 1 | 104.79 | 76.35667 | 181.1467 | 15.3993 | 31.61021 | 46.85822 |
| 2016 | 3 | 276.5567 | 1350.32 | 1626.877 | 36.26867 | 7.136617 | 43.04111 |
| 2016 | 4 | 175.58 | 1433.59 | 1609.17 | NA | NA | NA |
| 2016 | 5 | 38.20667 | 361.2 | 399.4067 | 6.825001 | 11.44844 | 17.74899 |
| 2016 | 7 | 44.10667 | 285.28 | 329.3867 | 2.17086 | 42.98332 | 42.41468 |
| 2016 | 9 | 124.5133 | 710.1967 | 834.71 | 30.24171 | 105.4086 | 76.82269 |
| 2016 | 11 | 152.69 | 238.27 | 289.1667 | NA | 25.29497 | 66.36763 |
| 2016 | 12 | 70.66 | 106.9267 | 177.5867 | 8.333925 | 18.41769 | 26.63488 |
| 2016 | 14 | 385.41 | 137.5533 | 522.9633 | 49.11438 | 14.57516 | 42.55046 |
| 2016 | 16 | 313.23 | 47.31333 | 360.5433 | 23.72018 | 5.411851 | 25.37919 |
| 2016 | 18 | 272.04 | 70.41667 | 342.4567 | 17.9971 | 22.97238 | 18.07291 |
| 2016 | 20 | 108.0133 | 55.61333 | 163.6267 | 8.421546 | 3.769421 | 4.990394 |
| 2016 | 22 | 129.8633 | 131.64 | 261.5033 | 25.61031 | 22.77144 | 47.29522 |
| 2016 | 24 | 147.0733 | 113.4967 | 260.57 | 6.511777 | 23.69947 | 30.02869 |
| 2016 | 26 | 45.51667 | 14.49333 | 60.01 | 4.736521 | 1.128952 | 5.528173 |

**Supplementary Table 4.**
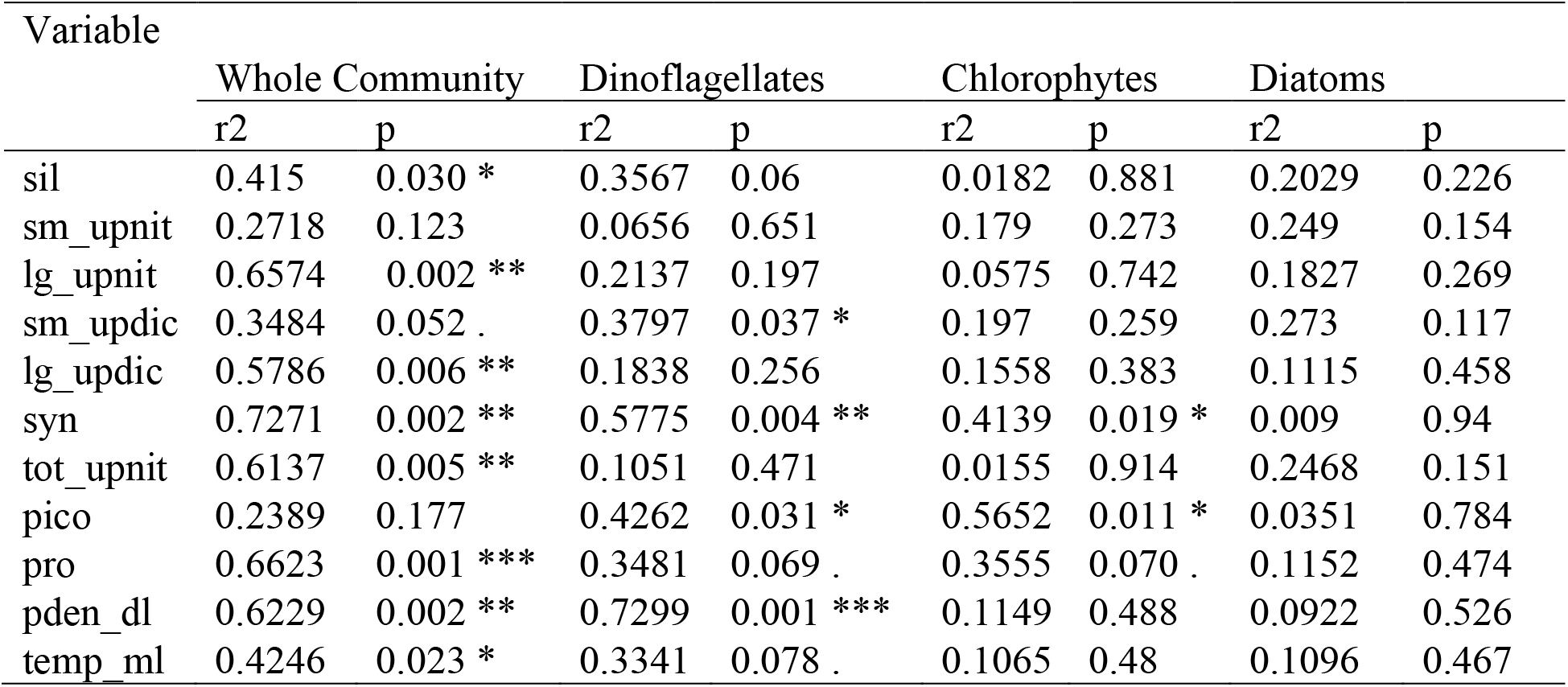
Results of the envfit() function. See supplementary Figure 2 for abbreviated variable key.

| Variable | Whole Community |  | Dinoflagellates |  | Chlorophytes |  | Diatoms |  |
| --- | --- | --- | --- | --- | --- | --- | --- | --- |
|  | r2 | p | r2 | p | r2 | p | r2 | p |
| sil | 0.415 | 0.030 * | 0.3567 | 0.06 | 0.0182 | 0.881 | 0.2029 | 0.226 |
| sm_upnit | 0.2718 | 0.123 | 0.0656 | 0.651 | 0.179 | 0.273 | 0.249 | 0.154 |
| lg_upnit | 0.6574 | 0.002 ** | 0.2137 | 0.197 | 0.0575 | 0.742 | 0.1827 | 0.269 |
| sm_updic | 0.3484 | 0.052 . | 0.3797 | 0.037 * | 0.197 | 0.259 | 0.273 | 0.117 |
| lg_updic | 0.5786 | 0.006 ** | 0.1838 | 0.256 | 0.1558 | 0.383 | 0.1115 | 0.458 |
| syn | 0.7271 | 0.002 ** | 0.5775 | 0.004 ** | 0.4139 | 0.019 * | 0.009 | 0.94 |
| tot_upnit | 0.6137 | 0.005 ** | 0.1051 | 0.471 | 0.0155 | 0.914 | 0.2468 | 0.151 |
| pico | 0.2389 | 0.177 | 0.4262 | 0.031 * | 0.5652 | 0.011 * | 0.0351 | 0.784 |
| pro | 0.6623 | 0.001 *** | 0.3481 | 0.069 . | 0.3555 | 0.070 . | 0.1152 | 0.474 |
| pden_dl | 0.6229 | 0.002 ** | 0.7299 | 0.001 *** | 0.1149 | 0.488 | 0.0922 | 0.526 |
| temp_ml | 0.4246 | 0.023 * | 0.3341 | 0.078 . | 0.1065 | 0.48 | 0.1096 | 0.467 |

**Supplementary Table 5.** Table with the values of depth, temperature, salinity, and density of the mixed (ML) and subthermocline (DL) layers obtained from CTD profiles.

| site | Year | Depth<br>(m) | | Potential<br>Density<br>$\rho_\theta$<br>(kg/m<sup>3</sup>) | | Salinity<br>(PSU) | | Temperature<br>(°C) | |
| --- | --- | --- | --- | --- | --- | --- | --- | --- | --- |
|  |  | DL | ML | DL | ML | DL | ML | DL | ML |
| 1 | 2015 | 74.574 | 41.765 | 1025.542 | 1023.962 | 35.07884 | 35.00383 | 17.15188 | 22.91208 |
| 3 | 2015 | 54.691 | 27.844 | 1025.282 | 1023.797 | 35.15576 | 34.94609 | 18.45271 | 23.33218 |
| 4 | 2015 | 58.667 | 4.972 | 1025.361 | 1022.756 | 35.15721 | 34.40935 | 18.14003 | 25.45745 |
| 5 | 2015 | 57.673 | 26.85 | 1025.231 | 1023.143 | 35.09401 | 34.58523 | 18.46886 | 24.62226 |
| 7 | 2015 | 44.748 | 21.878 | 1025.208 | 1022.855 | 35.11656 | 34.4859 | 18.62759 | 25.32338 |
| 9 | 2015 | 65.627 | 35.799 | 1025.274 | 1023.392 | 35.00837 | 34.77897 | 18.0309 | 24.28077 |
| 11 | 2015 | 57.673 | 10.939 | 1024.982 | 1021.94 | 35.00771 | 33.71368 | 19.18724 | 26.39509 |
| 12 | 2015 | 58.667 | 29.833 | 1024.497 | 1023.276 | 34.96258 | 34.70075 | 20.89355 | 24.46823 |
| 14 | 2015 | 65.627 | 18.895 | 1024.493 | 1022.858 | 34.79039 | 34.51659 | 20.42436 | 25.38764 |
| 16 | 2015 | 61.65 | 51.708 | 1025.428 | 1023.722 | 35.05016 | 34.91046 | 17.53284 | 23.50124 |
| 18 | 2015 | 70.598 | 42.759 | 1025.502 | 1023.586 | 35.04397 | 34.87231 | 17.20571 | 23.8639 |
| 20 | 2015 | 69.604 | 38.782 | 1025.196 | 1023.289 | 35.0318 | 34.70571 | 18.41766 | 24.43836 |
| 22 | 2015 | 62.644 | 20.884 | 1025.248 | 1023.283 | 35.07092 | 34.6674 | 18.32925 | 24.35941 |
| 24 | 2015 | 48.725 | 0.995 | 1025.424 | 1022.598 | 35.10714 | 33.9286 | 17.72452 | 24.7792 |
| 26 | 2015 | 65.627 | 26.85 | 1025.393 | 1022.549 | 35.11794 | 34.16701 | 17.88783 | 25.53792 |
| 1 | 2016 | 52.702 | 8.95 | 1026.097 | 1023.58 | 34.97879 | 34.21174 | 14.35497 | 22.14076 |
| 3 | 2016 | 25.856 | 6.961 | 1026.06 | 1025.435 | 34.9931 | 34.93188 | 14.57929 | 17.1086 |
| 4 | 2016 | 28.839 | 6.961 | 1026.049 | 1025.219 | 35.00383 | 34.95251 | 14.67041 | 18.06521 |
| 5 | 2016 | 49.719 | 1.989 | 1025.933 | 1024.737 | 34.99287 | 34.699 | 15.15999 | 19.21015 |
| 7 | 2016 | 27.844 | 13.923 | 1025.944 | 1024.843 | 35.0016 | 34.86895 | 15.14143 | 19.30295 |
| 9 | 2016 | 51.708 | 19.889 | 1025.828 | 1023.652 | 34.97766 | 34.28081 | 15.5809 | 22.07181 |
| 11 | 2016 | 47.731 | 16.906 | 1025.993 | 1024.203 | 34.9684 | 34.42807 | 14.79943 | 20.46199 |
| 12 | 2016 | 42.759 | 7.956 | 1026.054 | 1024.023 | 34.97158 | 34.33695 | 14.52727 | 20.87326 |
| 14 | 2016 | 26.85 | 22.872 | 1024.625 | 1024.115 | 34.88242 | 34.47773 | 20.18936 | 20.93247 |
| 16 | 2016 | 57.673 | 11.934 | 1026.055 | 1024.434 | 34.967 | 34.58984 | 14.5104 | 20.05481 |
| 18 | 2016 | 60.656 | 9.945 | 1026.007 | 1024.358 | 34.97825 | 34.58253 | 14.77365 | 20.32207 |
| 20 | 2016 | 46.736 | 38.782 | 1025.991 | 1023.382 | 34.98953 | 34.00239 | 14.88439 | 22.28094 |
| 22 | 2016 | 31.822 | 14.917 | 1025.946 | 1023.796 | 35.00014 | 34.24421 | 15.12943 | 21.45189 |
| 24 | 2016 | 45.742 | 10.939 | 1026.012 | 1023.403 | 35.01254 | 33.97096 | 14.86899 | 22.11866 |
| 26 | 2016 | 69.604 | 32.816 | 1025.977 | 1022.714 | 34.98422 | 33.64359 | 14.93502 | 23.66572 |

## Acknowledgments

We thank scientists and staff from the Galápagos Science Center (GSC), Universidad San Francisco de Quito (USFQ), and Galápagos National Park (GNP) for their logistical support during the cruises. We are grateful to Steve Walsh (UNC), Carlos Mena (USFQ) and Phil Page (UNC) for their efforts in coordinating the Galápagos Marine Expeditions. Others include Juan Pablo Muñoz (GSC), Leandro Vaca (GSC), Eduardo Espinoza (GNP), Jennifer Suarez (GNP) and the crew of the M/V Sierra Negra. We thank Sara Haines for assistance with processing and visualizing physical and chemical observations. We are grateful to Kimberly DeLong, Sharla Sugierski and Wilton Burns for assistance with sample analyses. Natalie Cohen, Rob Lampe, and Se Hyeon Jang provided helpful comments on the manuscript. Funding for this project was provided to A.M., H.S. and S.G from the Center for Galápagos Studies (Office of the Vice Chancellor for Research) and the UNC College of Arts and Sciences, to E.N. by the Andrew Marion Blackmon family trust and a National Science Foundation grant to A.M. (OCE1751805).

